# Cocaine addiction-like behaviors are associated with long-term changes in gene regulation, energy metabolism, and GABAergic inhibition within the amygdala

**DOI:** 10.1101/2022.09.08.506493

**Authors:** Jessica L. Zhou, Giordano de Guglielmo, Aaron J. Ho, Marsida Kallupi, Narayan Pokhrel, Hai-Ri Li, Apurva S. Chitre, Daniel Munro, Pejman Mohammadi, Lieselot LG Carrette, Olivier George, Abraham A. Palmer, Graham McVicker, Francesca Telese

## Abstract

The amygdala processes positive and negative valence and contributes to the development of addiction, but the underlying cell type-specific gene regulatory programs are unknown. We generated an atlas of single nucleus gene expression and chromatin accessibility in the amygdala of outbred rats with low and high cocaine addiction-like behaviors following prolonged abstinence. Between rats with different addiction indexes, we identified thousands of cell type-specific differentially expressed genes enriched for energy metabolism-related pathways that are known to affect synaptic transmission and action potentials. Rats with high addiction-like behaviors showed enhanced GABAergic transmission in the amygdala, which, along with relapse-like behaviors, were reversed by inhibition of Glyoxalase 1, which metabolizes the GABA_A_ receptor agonist methylglyoxal. Finally, we identified thousands of cell type-specific chromatin accessible sites and transcription factor (TF) motifs where accessibility was associated with addiction index, most notably at motifs for pioneer TFs in the Fox, Sox, helix-loop-helix, and AP1 families.

## Introduction

The amygdala is a key brain region involved in regulating a wide range of behaviors, including those related to emotions, motivation and memory^1^. In response to rewarding or aversive environmental stimuli, the amygdala allows organisms to engage in subsequent valence-specific behaviors by determining the value of different stimuli and guiding decision-making based on potential outcomes^1^. The amygdala is implicated in numerous neuropsychiatric disorders including addiction^2–4^. Repeated drug use leads to a heightened sense of pleasure, which engages the amygdala to form drug-associated memories and reinforces drug-seeking behavior, as the individual is motivated to seek out and use the drug again in order to experience the rewarding effects^5^. In addition, during withdrawal from addictive drugs, the amygdala mediates negative emotional states, such as anxiety, fear, and irritability^5^. Avoidance of these aversive emotions enhances the incentive value of the drug, leading to sustained drug-seeking behaviors and relapse^6–8^. Because prevention of relapse is the cornerstone of effective treatments for addiction, it is important to understand the amygdala’s role in addiction and relapse.

The amygdala is composed of multiple discrete and interconnected subregions, each characterized by highly specialized neuronal populations distinguishable by their morphology and electrophysiological properties^9^. The major subdivisions include the basolateral amygdala (BLA), composed of excitatory glutamatergic neurons and GABAergic inhibitory interneurons, and the central amygdala (CeA), composed of GABAergic neurons^10–12^. While the behavioral function and connectivity of individual subregions of the amygdala have recently been established^1^, the mechanisms by which distinct subpopulations of neuronal and non-neuronal cells contribute to its function remains unclear.

Single-cell genomics is a powerful new approach for determining the cellular function and diversity of complex tissues like the amygdala. Single-cell RNA-sequencing (scRNA-seq), which profiles gene expression in individual cells, has identified and cataloged diverse cell types in human, mouse, and non-human primate brains^13–19^. In addition, single-cell assays for transposase-accessible chromatin (scATAC-seq), which profile chromatin accessibility at single cell resolution, have identified regulatory DNA sequences in the rodent and human brain^13,20–26^. Regulatory elements identified by scATAC-seq include promoters and enhancers, which confer cell type-specificity to gene expression by recruiting sequence-specific transcription factors (TFs)^27–30^.

Single cell assays have the potential to reveal, at a molecular level, how specialized amygdalar cell populations are involved in addiction. For example, given that most genetic variants associated with complex human diseases like addiction are located in noncoding regions of the genome^31^, snATAC-seq could uncover genetically determined, cell-type specific differences and facilitate functional interpretation of genetic variants^32^. Thus far, however, the application of single-cell assays to the study of addiction-like behaviors in rodents has been limited. Single nucleus RNA-seq (snRNA-seq) has been applied to characterize cellular diversity in brain regions involved in the reward system^33–36^, and has been used to analyze transcriptional changes induced by cocaine and morphine^37,38^. However, these prior studies used isogenic rodents, which means that genetically-mediated differences in susceptibility to addiction-like behaviors were not examined. Furthermore, these studies performed experiments following acute, experimenter-administration of drug treatments, which means that they reflect the acute effects of involuntary drug use rather than molecular differences associated with the development of long-lasting addictive-like behaviors. For these reasons, the results from prior single nucleus studies have significant limitations.

To address this knowledge gap, we performed snRNA-seq and snATAC-seq using amygdala tissue from outbred rats obtained from a large genetic study of cocaine addiction-related traits^39^. These rats are subjected to prolonged abstinence from voluntary cocaine intake in a well-validated model of extended access to drug intravenous self-administration (IVSA)^6,39–41^. IVSA is associated with neurochemical changes in key brain regions, which are thought to be similar to those observed in humans with cocaine use disorder^42^. This study used outbred heterogeneous stock (HS) rats because they have high levels of genetic variation and rich phenotypic diversity^43–46^. By analyzing differences in gene expression and chromatin accessibility in rats with high and low addiction indexes (AI), we identified genes and transcriptional regulators associated with cocaine addiction-like behaviors, including those implicated in energy metabolism and neurotransmitter pathways. Furthermore, using genetic predictions of gene expression, we found that genetic differences contribute to the gene expression differences between high and low AI rats. Finally, we performed pharmacological manipulation of GABA_A_ receptor signaling in amygdalar tissue slices and in rats to validate insights gained from the transcriptomic data.

## Results

### Behavioral characterization of HS rats exhibiting low or high cocaine addiction-like traits

To investigate how chronic cocaine use influences cellular states associated with addiction-like behaviors, we performed snRNA-seq and snATAC-seq on amygdala tissues from HS rats subjected to protracted abstinence (4 weeks) following extended access to cocaine IVSA ^39,47–50^ (Fig. 1a). The animals were trained to self-administer cocaine via lever press (0.5 mg/kg/infusion) in operant chambers in short access (ShA, 2h/day, 5 days per week) and long access (LgA, 6h/day, 5 days/week) sessions. We measured the number of cocaine infusions, or lever presses, during each session of the behavioral protocol to quantify escalation of intake, motivation, and compulsive-like behavior. Specifically, we measured escalation as the increase in the mean number of cocaine rewards during each of the LgA sessions compared to the first day of the LgA phase; we measured motivation as the mean number of cocaine rewards over the ShA and LgA sessions under a progressive ratio (PR) schedule of reinforcement, which is when the number of lever presses required to obtain a cocaine infusion was progressively increased; and we measured compulsive-like behavior as drug taking despite adverse consequences, by pairing 30% of lever presses with an electric foot shock (Fig. 1b). Based on individual behavioral measures for each rat (Fig. 1c), we calculated an addiction index (AI)^39^ as the average of the normalized values (z-scores) of the three behavioral measures. Prior work has demonstrated that AI measures vulnerability (high AI) or resilience (low AI) to developing cocaine addiction-like behaviors^39,51–53^.

**Figure 1.**
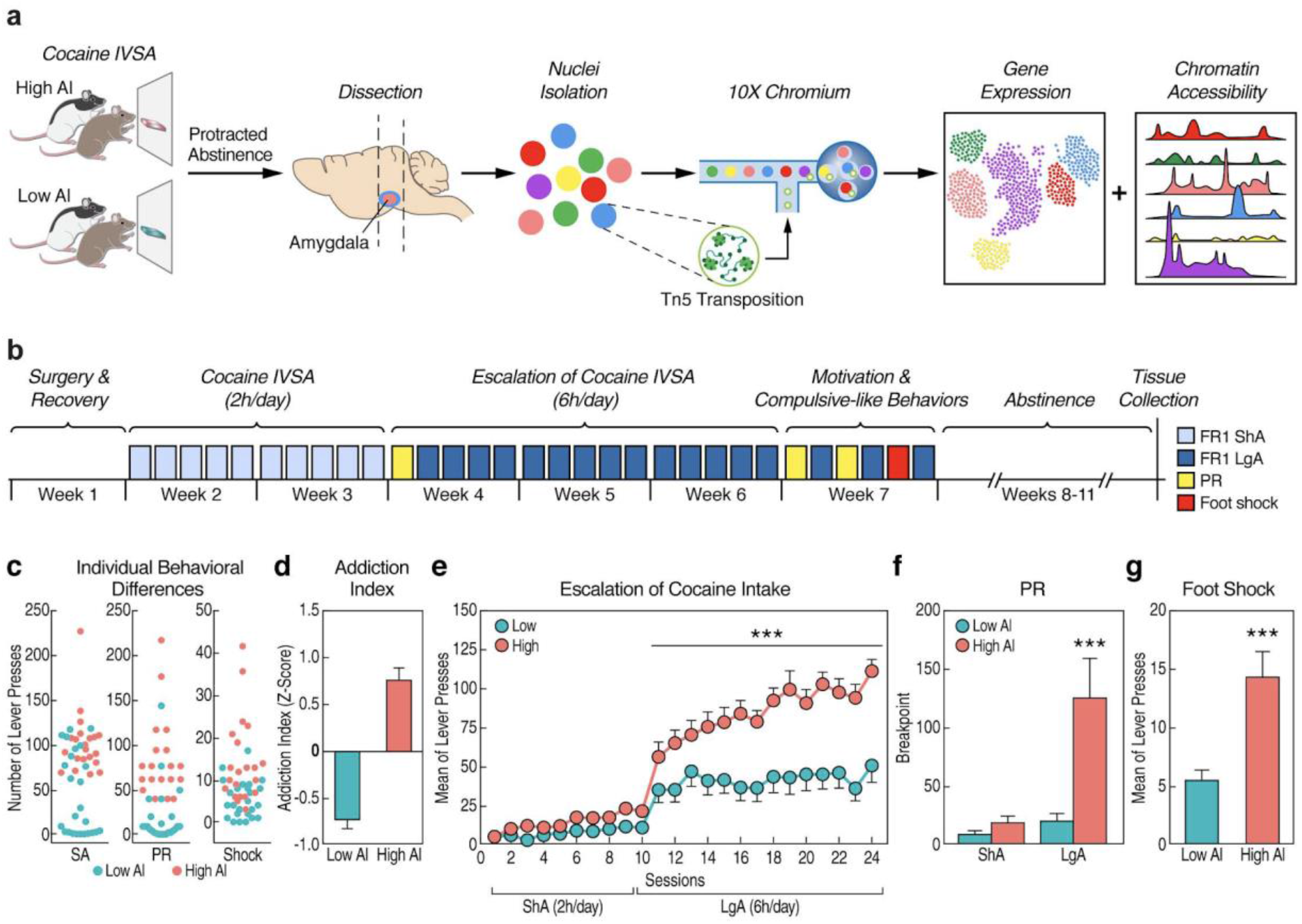
Experimental design and rat IVSA cocaine model of addiction. **a)** Schematic of the study design. **b)** Timeline of the behavioral protocol. **c)** Individual differences in total number of lever presses in self-administration (SA), progressive ratio (PR) and shock-paired (Shock) sessions for each rat. **d)** Mean addiction index scores in high and low AI rats. **e)** Mean number of lever presses across each ShA and LgA IVSA session in high (n=21) and low (n=25) AI rats (*** p < 0.001, two-way repeated measures ANOVA interaction time x group F_13,572_=4.175). **f)** Breakpoint analysis of high (n=21) and low (n=25) AI rats under ShA versus LgA (*** p<0.001 mixed effect model, addiction index × phase interaction, p=0.0049, F_1,41_=8.83). **g)** Mean number of lever presses when paired with electric footshock in high (n=21) and low AI (n=25) rats (***p<0.001, unpaired Student’s t-test, t_44_=3.936). Error bars in panels d-g represent the standard error of the mean.

We classified rats into high and low AI groups (Fig. 1d). Both high and low AI rats acquired fewer cocaine rewards in the ShA compared to the LgA sessions of the IVSA protocol (Fig. 1e, two-way repeated measures ANOVA, addiction index × phase interaction p<0.0001, F_23,1012_=8.523). There was no difference between groups in cocaine rewards during ShA sessions. However, we observed a contrasting pattern in escalation during LgA sessions. During LgA sessions, rats with high AI exhibited a progressive escalation of drug intake compared to rats with low AI as evidenced by their increased number of cocaine infusions over the course of this phase of the IVSA protocol (two-way repeated measures ANOVA interaction time x group F_13,572_=4.175, p < 0.0001, Fig. 1e). In contrast, low AI rats did not show escalation during the LgA sessions (Fig. 1e).

During PR sessions, motivation for cocaine increased in the high AI rats but not in low Al rats when comparing ShA versus LgA (Fig. 1f, mixed effect model, addiction index × phase interaction, p=0.0049, F_1,41_=8.83; Bonferroni corrected p=0.0001, post hoc comparisons). Finally, high AI rats showed increased responses despite adverse consequences compared to low AI rats, as demonstrated by the higher number of cocaine infusions when the reward was paired with an electric foot shock (Fig. 1g, p<0.001, unpaired Student’s t-test, t_44_=3.936), which may reflect compulsive-like drug use. These results show that we can capture multiple behavioral aspects that are relevant to cocaine use disorders by using this model of extended access to cocaine IVSA in outbred rats.

### snRNA-seq and snATAC-seq defines distinct populations of cell types in the amygdala

The amygdala is thought to contribute to the development of addiction through its regulation of drug-seeking behavior, which, in rats, progressively increases after withdrawal from drug IVSA^6,54^. To identify neuroadaptations that persist in the amygdala after chronic drug exposure during the withdrawal stage, we collected amygdalae after 4 weeks of abstinence from cocaine IVSA (Fig. 1a). We purified nuclei and measured the gene expression and chromatin accessibility profiles of individual nuclei by performing snRNA-seq and snATAC-seq with the 10X Genomics Chromium workflow. We performed these experiments on high and low AI rats, as well as naive rats never exposed to cocaine. For snRNA-seq, we used 19 rats including 6 with high AI, 6 with low AI, and 7 naive rats (Supplementary Data 1). For snATAC-seq we used 12 rats including 4 with high AI, 4 with low AI, and 4 naive rats (Supplementary Data 2).

After filtering low quality nuclei and potential doublets based on quality metrics, we obtained a combined total of 163,003 and 81,912 high quality nuclei from the snRNA-seq and snATAC-seq samples, respectively (Fig. S1-6, Supplementary Data 3-4). Across the snRNA-seq samples, the mean reads per cell varied from 11,967 to 50,343 and the median number of detected genes ranged from 1,293 to 2,855. Across the 12 snATAC-seq samples, the median number of high-quality fragments per nucleus ranged from 7,111 to 22,018. Across samples, we observed means of 8579 and 6826 nuclei per rat in the snRNA-seq and snATAC-seq datasets, respectively (Fig. S7). The above metrics are consistent with previously published single-nucleus sequencing studies of the amygdala^33,55^. Using these data, we performed normalization, integration across rats, dimensionality reduction and clustering using Seurat^56^ (for snRNA-seq) and Signac^57^ (for snATAC-seq). In total, we identified 49 cell type clusters in the integrated snRNA-seq dataset and 41 cell type clusters in the integrated snATAC-seq dataset (Fig. S8). Visualization of the integrated data indicated that the clustering is not influenced by batch effects such as sequencing library, percentage of mitochondrial DNA, or individual rats^58^ (Fig. S9).

We annotated the snRNA-seq clusters based on the expression of established cell type-specific marker genes^59–63^ (Fig. 2a-b). The major cell types included excitatory neurons (denoted by expression of *Slc17a7)*, inhibitory GABAergic neurons (*Gad1/Gad2)*, astrocytes (*Gja1)*, microglia (*Ctss)*, mature oligodendrocytes (*Cnp)*, oligodendrocyte precursor cells (OPC) (*Pdgfra)*, and endothelial cells (*Cldn5)* (Fig. 2c). To annotate the snATAC-seq clusters, we estimated gene activity from pseudo bulk chromatin accessibility at promoter regions of cell marker genes and used these gene activity scores to impute gene expression in the snATAC-seq samples (Fig. S10). The imputed gene expression clearly delineates the cell clusters into the same major cell types described above demonstrating strong concordance between our snRNA-seq and snATAC-seq data (Fig. 2d). In addition to the major cell types, we also identified seven subtypes of inhibitory neurons based on the expression of known cell marker genes (Fig. 2e). Cell type proportions appeared to be consistent across samples (Fig. S11-12). We also sub-clustered the excitatory neurons and identified 18 distinct clusters (Fig. S13), with top markers including known subpopulation markers such as *Cdh13, Nr4a2, Bdnf*^64^.

**Figure 2.**
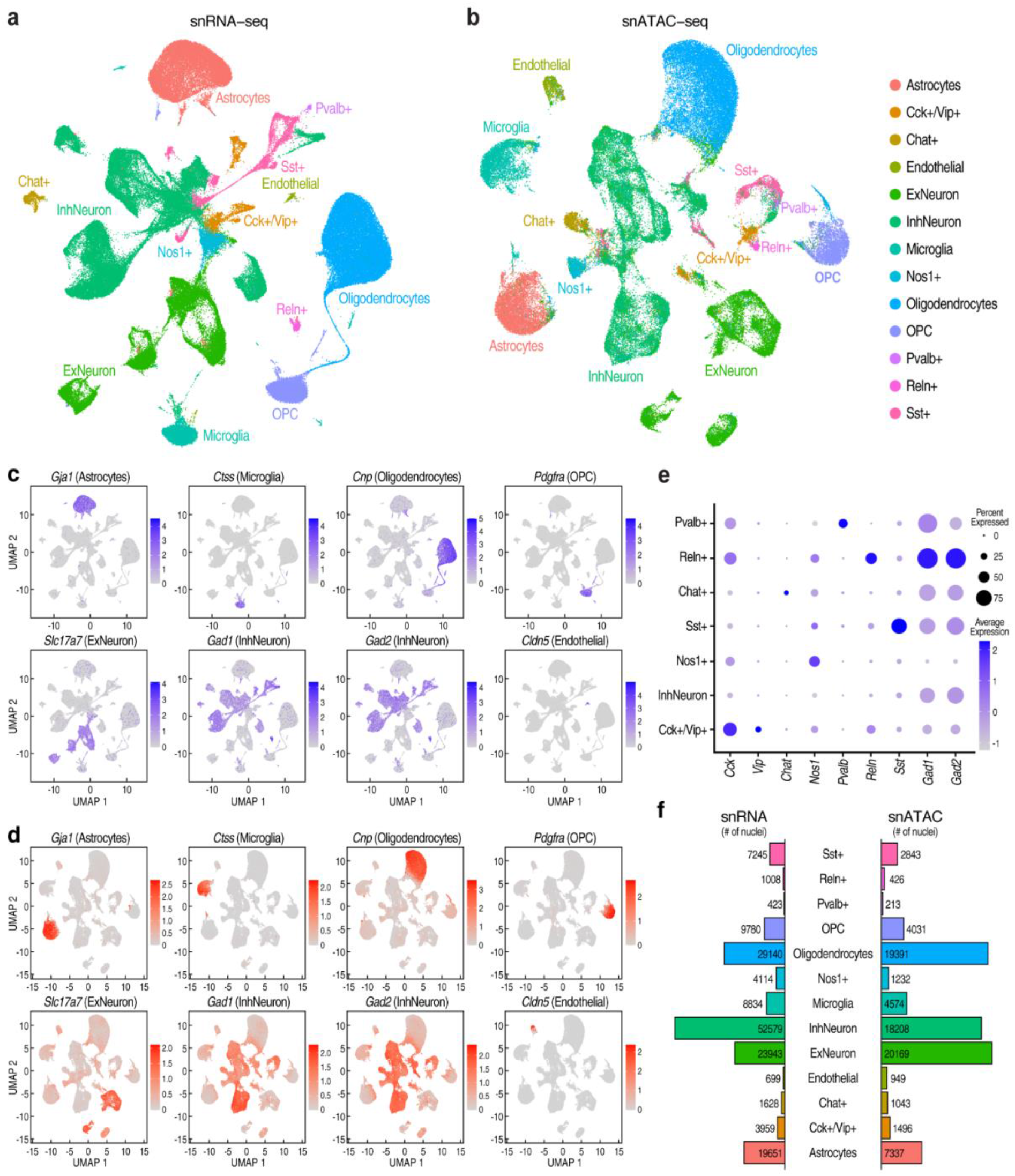
Summary of single nucleus RNA-seq and ATAC-seq data from the rat amygdala. **a)** Uniform Manifold Approximation And Projection (UMAP) plot of single nucleus RNA-seq (snRNA-seq) data from the rat amygdala. Data are combined across 19 samples, with high, low, and naive addiction index labels. Cells are colored by cluster assignments performed with K-nearest neighbors. We assigned cell type labels to the clusters based on the expression of known marker genes. **b)** UMAP plot of single nucleus ATAC-seq data from 12 rat amygdala samples. snATAC-seq data was integrated with the snRNA-seq data and cluster labels were transferred to the snATAC-seq cells. **c)** Feature plot showing expression of marker genes used to label major subsets of cells: *Gja1* (astrocytes), *Ctss* (microglia), Cnp (oligodendrocytes), *Pdgfra* (oligodendrocyte precursor cells (OPCs), *Slc17a7* (excitatory neurons), *Gad1* and *Gad2* (inhibitory neurons), and *Cldn5* (endothelial cells). **d)** Feature plot showing imputed gene expression of cell type-specific marker genes in snATAC-seq dataset. **e)** Expression of marker genes in cell clusters corresponding to highly specific subsets of inhibitory neurons. The shading and diameter of each circle indicate the estimated mean expression and the percentage of cells within the cluster in which the marker gene was detected. **f)** The number of nuclei assigned to each cell type cluster for the snATAC-seq and snRNA-seq datasets.

The total number of nuclei we obtained for each cell type varied substantially (Fig. 2f). As expected, excitatory and inhibitory neurons are the most common major cell types. The snRNA-seq dataset contains 52,579 (∼32.3%) nuclei from inhibitory neurons and 23,943 (∼14.7%) nuclei from excitatory neurons (Table S1). The snATAC-seq dataset contains 18,208 (∼22.2%) nuclei from inhibitory neurons and 20,169 (∼24.6%) nuclei from excitatory neurons (Table S1). Endothelial cells and some subtypes of inhibitory and excitatory neurons have small numbers of nuclei in the dataset, so for most downstream analyses we focused on the six most common major cell types (Fig. 2a-b).

To determine how the cell types we identified in the whole amygdala correspond to cell types within spatially defined amygdalar subregions, we generated snRNA-seq data from the CeA and BLA (Fig. S14). We found that cell clusters from the CeA and BLA were distinct from one another, but these regions collectively contained most cell types also identified in the whole amygdala (Fig. S14a). Consistent with the known cell type composition of the CeA and BLA^65^, the cell clusters from the CeA co-clustered primarily with inhibitory neurons while the cell clusters from the BLA co-clustered with excitatory neurons (Fig. S14b). Glial cell types from the whole amygdala contained cells from both subregions, with the exception of astrocytes, which mostly co-clustered with cells from the CeA but not the BLA, suggesting that astrocytes might play a specific role in BLA-related function (Fig. S14a-b).

In combination, the snRNA-seq and snATAC-seq datasets that we generated are the first single-cell atlas of molecularly defined cell types in the rat amygdala. The inclusion of multiple high AI, low AI, and naive rats make these datasets an important resource for studying gene expression and chromatin accessibility in the amygdala under both normal conditions as well as in the context of cocaine addiction-like behaviors.

### Measuring cell type-specific differential gene expression between rats displaying a high versus a low addiction index for cocaine

We used the negative binomial test to identify differentially expressed genes (DEGs) between high and low AI rats in each cell type (Fig. 3a-b, Supplementary Data 5). To control for batch effects or violations in the differential expression model assumptions (for example, unaccounted for overdispersion in the data) that can cause deflated (overly significant) p-values, thereby yielding false signals of differential expression^66,67^, we performed the same statistical test after permuting the AI labels of the rats. This permutation simulates a null distribution where there is no association between AI and gene expression. This approach is often used to assess p-value calibration in QTL studies^68,69^. While the results from the unpermuted data are highly enriched for low p-values, the p-values from the permuted data resemble the null expectation. This indicates that the highly-significant DEGs we identified are not due to poor p-value calibration or batch effects (Fig. S15, Supplementary Data 6).

**Figure 3.**
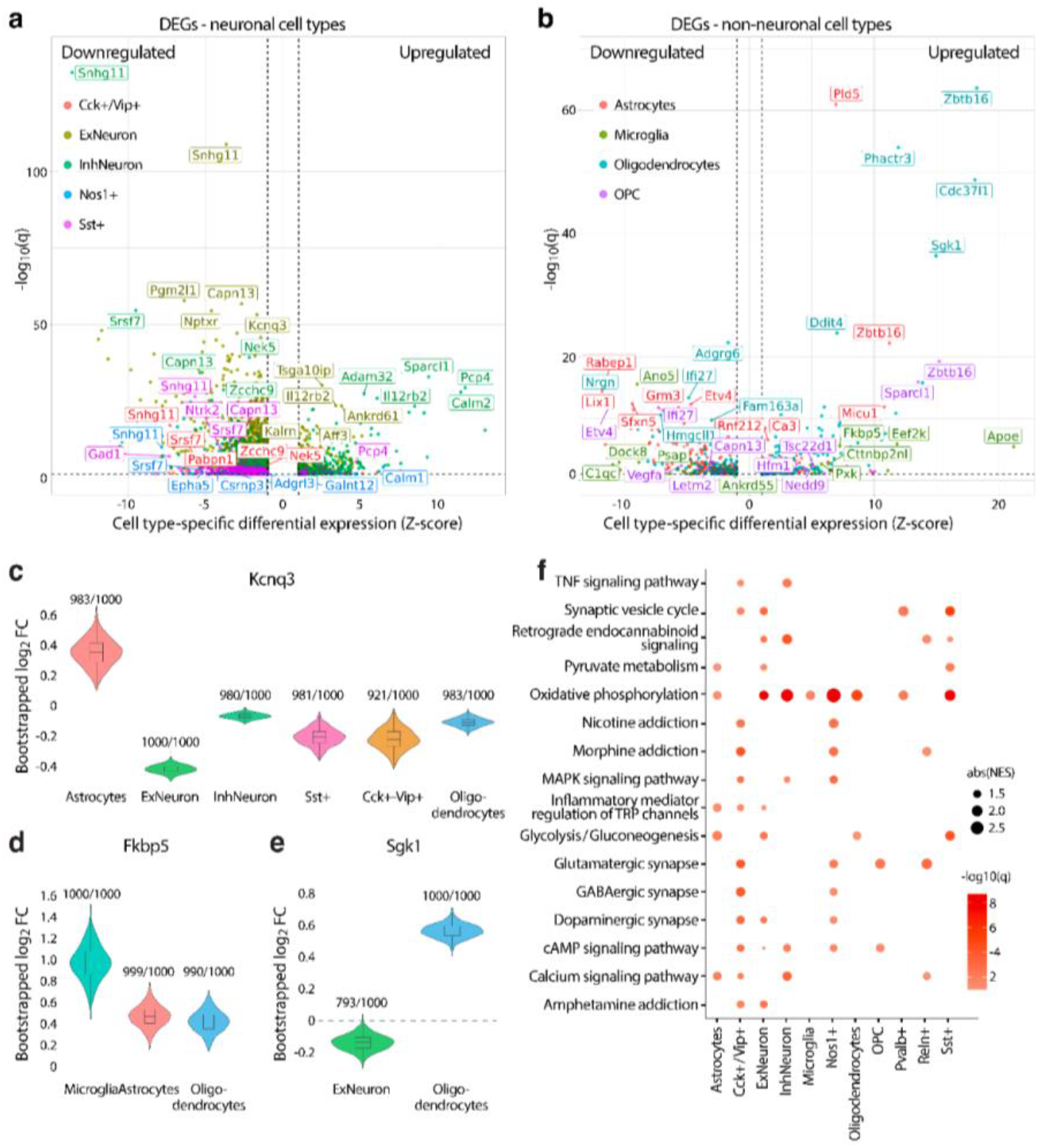
Differential gene expression between high and low addiction index rats. **a)** Volcano plot summarizing differential gene expression between high and low AI rats. Points are colored by cell type, and the five most significant (FDR<10%) up- and downregulated genes in each cell type are indicated with labels. Within each cell type, we normalized the log fold changes reported by Seurat with a z-score and plotted the cell type-specific z-scores of the log fold changes on the x-axis (positive fold change = higher expression in high AI rats; negative fold change = higher expression in low AI rats). The -log_10_ false discovery rate (FDR) corrected p-values (q-values) are plotted on the y-axis. **b)** Volcano plot summarizing differential gene expression between high and low AI rats for non-neuronal (glial) cell type clusters. **c-e)** Violin plots showing distribution of log_2_FC from the negative binomial test performed in 1000 bootstrap iterations. Fractions indicate the number of bootstrap iterations in which the log2FC estimate was significantly different than 0. Bootstrap distributions were obtained for cell types in which the following genes had significant differential expression (FDR<10%): **c)** *Kcnq3*; **d)** *Fkbp5*; **e)** *Sgk1*. **f)** KEGG pathways that are enriched for differentially expressed genes by cell type. Size of dot indicates -log10(q) while color indicates normalized enrichment score (NES), which is a metric of GSEA. Only pathways/cell types where q<0.1 are visualized.

We grouped DEGs into small (abs(log_2_FC)<0.1) or large effect size groups (abs(log_2_FC)≥0.1) and observed that most of the significant DEGs (FDR<10%) have small effect sizes (Fig. S16). In total, we identified 557 unique significant DEGs with large effect sizes in at least one cell type and 8,775 unique significant DEGs with small effect sizes in at least one cell type. The number of significant DEGs between high and low AI rats correlates with the size of the cell type population (Fig. S16), which likely reflects greater power to detect differential expression in common cell types. Most of the large effect DEGs (431, or 75%) are also small-effect DEGs in at least one other cell type, indicating that while there are shared patterns of differential expression across cell types, the effect sizes vary across cell types. We also found that significant DEGs were enriched for gene expression quantitative trait loci (eQTLs), which are genetic variants associated with a gene’s expression, in rat brain tissues^70^ in almost every cell type tested (Chi-squared test with 1 degree of freedom, p<0.05) (Fig. S17, Table S2). This suggests that heritable differences influence the changes in expression that we observed. Among the most significant DEGs with eQTLs (Supplementary Data 7), we identified genes with previously reported roles in cocaine or other substance use disorders. For example, *Kcnq3* was differentially regulated across neuronal and glial cell types, and this gene encodes a subunit of a potassium channel implicated in the regulation of reward behavior and susceptibility to drug addiction (Fig. 3c)^71–73^. Additionally, *Fkbp5* and *Sgk1*, two transcriptional targets of the glucocorticoid receptor, were differentially regulated specifically in glial cell types, and these genes are associated with reward behavior and drug addiction vulnerability (Fig. 3d-e)^74–76^. These results suggest that genetic differences contribute to the gene expression differences between rats with high and low AI.

The observed DEGs could reflect pre-existing genetic differences or the differential exposure to cocaine in the high versus low AI groups. To further examine the contribution of genetics to observed differences in gene expression, we leveraged genotypes and gene expression data from a previously published reference population of 339 naive HS rats^70^. This allowed us to predict gene expression based on cis-genetic variation in the absence of cocaine exposure. Specifically, we trained models which predict gene expression from SNP genotypes^77^ using whole brain bulk RNA-seq from 339 naive HS rats. We then used these models to predict the expression of genes with at least one cis-acting eQTL (8,997 genes) in each of the rats in our snRNA-seq dataset. We compared the differences in mean predicted expression in the high versus low AI rats to the observed differences in expression for each cell type. Before correlating predicted expression to observed expression, we filtered out genes for which the predictive models had low Pearson *r*^2^, because genes with higher *r*^2^ have more of their variance in expression explained by cis-genetic variation (Table S3). Among our major cell types, the observed and predicted expression differences were significantly correlated (Spearman’s *ρ*, p<0.05) for microglia, oligodendrocytes and inhibitory neurons (Fig. S18, Table S3). We found that increasing the stringency of the *r*^2^ cutoff increased the strength of these correlations (Fig. S18, Table S3). These observations indicate that genetic differences in high versus low AI rats contribute to at least some of the observed differences in expression between high vs. low AI rats. Cocaine is also likely to contribute to the differences in expression; however, the relative contributions of cocaine and genetics are difficult to quantify due to limitations in the genetic predictions of gene expression.

To identify pathways with altered regulation between high and low AI rats, we performed gene set enrichment analysis (GSEA)^78,79^ of KEGG pathways using estimates of differential gene expression (log_2_ fold change) for each cell type. We identified significant enrichment of several pathways related to addiction (e.g. amphetamine, nicotine, and morphine addiction), neurotransmission (e.g. synaptic vesicle cycle, GABAergic synapse, glutamatergic synapse, and dopaminergic synapse), energy metabolism (e.g. glycolysis, pyruvate metabolism, and oxidative phosphorylation), and others (Fig. 3f, Supplementary Data 8). Most cell types showed enrichment of genes belonging to the oxidative phosphorylation pathway, which, together with glucose metabolism, is the main energy source for synaptic activity and action potentials^80,81^. Moreover, different subtypes of inhibitory neurons as well as excitatory neurons were enriched for synaptic vesicle cycle and synapse-related pathways. In combination, these observations suggest that addiction-like behaviors are associated with alterations in the metabolic state of amygdalar cell populations, which can directly impact neural network activity within the amygdala by affecting neurotransmission and synaptic pathways.

### The development of cocaine addiction-like behaviors is linked to elevated GABAergic transmission in the amygdala

To test the hypothesis that altered metabolic state in amygdalar cells changes neural activity within the amygdala, we focused on GABAergic transmission because alterations of this neurotransmitter system have been previously described in the amygdala in the context of addiction-related phenotypes^2^. Specifically, we measured GABAergic transmission by recording spontaneous inhibitory postsynaptic currents (sIPSCs) in the central amygdala (CeA). CeA slices were collected from a separate cohort of 5 low AI and 5 high AI HS rats that were subjected to prolonged abstinence following the same behavioral protocol described for the snRNA-seq and snATAC-seq experiments (Fig. 4a). As a control, we used CeA slices prepared from 5 age-matched naive HS rats to record baseline GABAergic transmission. There were differences in mean sIPSC frequencies among the groups (one-way ANOVA F_2,22_=6.77, p=0.0051), reflecting a progressive increase in GABAergic transmission from naive to low AI to high AI rats (Fig. 4b, Fig. S19a), without detectable changes in amplitude (Fig. S19b-c). These results are consistent with the hypothesis that the cocaine addiction-like behaviors exhibited by high AI rats alters GABAergic transmission.

**Figure 4.**
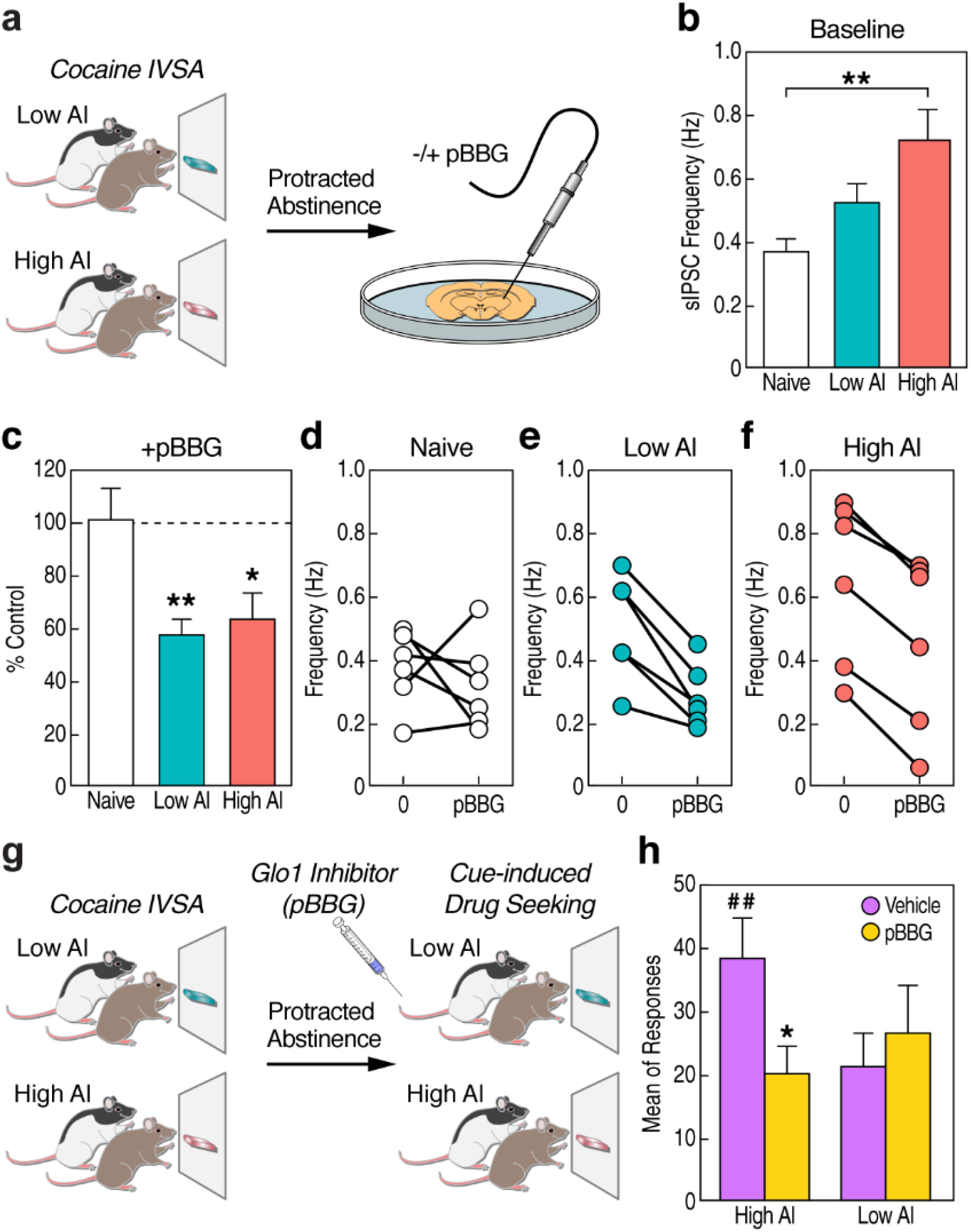
Electrophysiology and GLO1 inhibition experiments implicate GABAergic inhibition in cocaine addiction-like behaviors. **a)** Schematic showing animal model used for electrophysiology recording in CeA slices from HS rats subjected to 4 weeks of abstinence from cocaine IVSA. Electrophysiological recordings were taken before and after pBBG (S-bromobenzylglutathione cyclopentyl diester) treatment. **b)** Baseline iPSC frequency (measured before pBBG injection). Significant differences in the means between the three groups were observed (**p < 0.01; one-way ANOVA F_2,22_=6.77, followed by post-hoc comparison using Tukey’s HSD). **c)** sIPSC frequency following pBBG treatment. We observed reduced frequency in the CeA slices from high and low AI rats following pBBG treatment (**p < 0.01, *p<0.05 following Bonferroni correction; unpaired two-sided Student’s t-test). Change in sIPSC frequency following pBBG treatment in **d)** naive, **e)** low, and **f)** high rats**. g)** Schematic of animal model used to test cocaine-seeking behavior. Rats were injected with pBBG following a period of prolonged abstinence and re-exposed to the self-administration chambers in the absence of cocaine. **h)** Following injection of pBBG, high AI rats (n=12) showed significantly higher cocaine-seeking behavior compared to low AI rats (n=14), which was reduced by pBBG treatment (## p<0.001, *p<0.05 following Bonferroni correction; two-way ANOVA for each measure). Error bars in panels b,c, and h represent the standard error of the mean.

To further investigate the link between GABAergic transmission and energy metabolism in the amygdala with cocaine addiction-like behaviors, we measured siPSCs frequency and amplitude before and after application of S-bromobenzylglutathione cyclopentyl diester (pBBG)^82,83^. pBBG is a small molecule inhibitor of glyoxalase 1 (GLO1), the rate limiting enzyme for the metabolism of methylglyoxal (MG), which is a byproduct of glycolysis that acts as a competitive partial agonist of GABA_A_ receptors^82^. We found that pBBG reduced the sIPSC frequency compared to vehicle for both high and low AI rats (paired t-tests, t_5_=11.83, p=7.6e-5 and t_5_=5.07, p=3.9e-3, respectively), but not naive rats (t_5_=0.71, p=0.51) (Fig. 4c-f, Fig. S19a). We observed no effect on iPSCs amplitude (Fig. S19b-c). In most situations, changes in frequency of events indicate presynaptic modulation while changes in amplitude of events reflect postsynaptic modulation; however, previous studies have shown that GABA modulates synaptic transmission presynaptically^84,85^. These findings suggest that Glo1 inhibition may alter presynaptic GABA-A receptor function, leading to reduced GABA release at inhibitory terminals and suppression of inhibitory connections within the CeA.

These results led us to hypothesize that GLO1 inhibition would revert behavioral responses after prolonged abstinence from cocaine IVSA. To test this hypothesis, we measured cue-induced reinstatement of cocaine seeking behavior in a separate cohort of 26 low and high AI rats 30 minutes after systemic injection of pBBG or vehicle^86^ following 4 weeks of abstinence from cocaine IVSA (Fig. 4g). During this test, rats were subjected to the same operant conditions of cocaine IVSA, but without drug availability. Then, reinstatement was triggered by re-exposure to the cocaine infusion-associated light cue. The two-way repeated measures ANOVA showed a significant interaction between AI and pBBG treatment (F_1,24_=6.609, p<0.05), indicating that pBBG versus vehicle reduced cue-induced reinstatement in high AI rats (p-value<0.05, post hoc comparisons with Bonferroni correction), but not in low AI rats (p>0.05). Overall these results demonstrate that modulating GABA_A_ transmission via the pharmacological inhibition of GLO1 decreases relapse-like behaviors in animals with high cocaine AI.

### Mapping differences in chromatin accessibility associated with cocaine addiction-like behaviors

To identify regions of accessible chromatin from the snATAC-seq data, we used MACS2^87^ to call peaks from the aligned reads for each rat and created a union peak set across rats. We examined pseudo bulk chromatin accessibility at the transcription start site (TSS) of selected cell type marker genes and observed cell type-specific patterns of accessibility at the expected marker genes of each cell type (Fig. 5a, Fig. 2c-d), indicating that the chromatin accessibility corresponds well with the transcriptome measurements.

**Figure 5.**
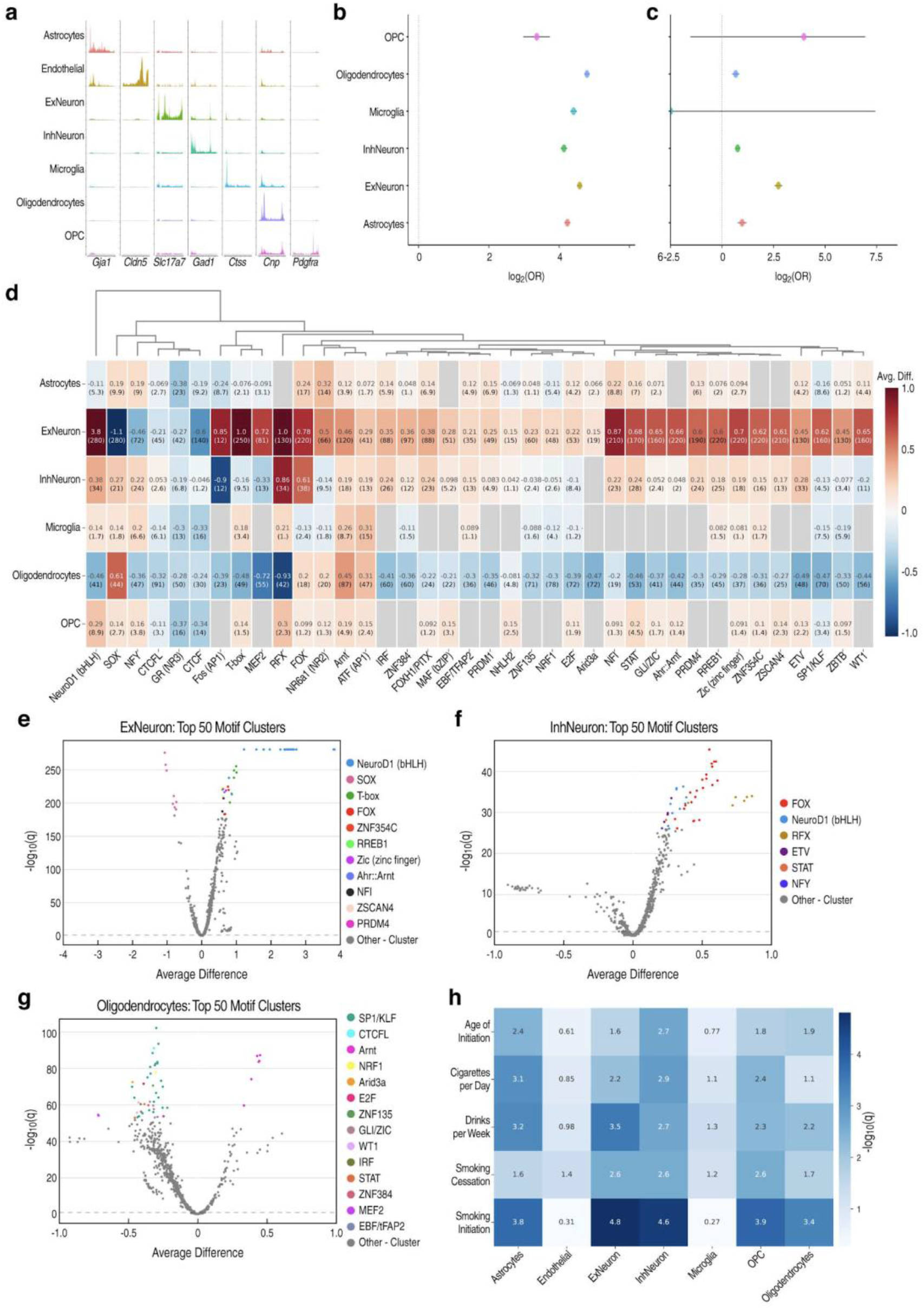
Analysis of chromatin accessibility and regulatory elements involved in cocaine dependence. **a)** Pseudobulk chromatin accessibility at the promoter regions of marker genes for major cell types identified in our analysis. **b)** Significant DEGs (FDR<10%) for each major cell type are enriched for promoters with differentially accessible chromatin accessibility (FDR<10%; Fisher’s exact test) in the snATAC-seq data. This indicates that the snRNA-seq and snATAC-seq results are consistent and indicate that long-term transcriptional changes are associated with changes in chromatin accessibility of gene promoters. **c)** Cell type-specific differentially accessible peaks (FDR<10%; Fisher’s exact test) are enriched in TSS/promoter regions compared to non-TSS/promoter regions. Error bars in b,c represent 95% confidence intervals for log2 odds ratios (ORs). **d)** Heatmap showing differential activity of various motifs in the significant differential peaks of each cell type. Values indicate average difference of chromVar deviation scores with -log10(p) in parentheses below. There are many cases where motifs display increased activity in the peaks which are more accessible in upregulated peaks in neurons while also displaying decreased activity in downregulated peaks in oligodendrocytes. **e-g)** Volcano plots showing average difference (x-axis) and - log10(q) (y-axis) of chromVAR deviation scores for top 50 motif clusters in **e)** excitatory neurons, **f)** inhibitory neurons, and **g)** oligodendrocytes. **h)** LD score regression results showing significance (-log10p) of enrichment of heritability for several traits related to alcohol and nicotine addiction in cell type-specific accessible chromatin regions (mapped to hg19).

To better understand the regulatory mechanisms involved in cocaine addiction, we analyzed differences in chromatin accessibility between high and low AI rats. We performed negative binomial^88,89^ tests to measure cell type-specific differential chromatin accessibility (Supplementary Data 9), and compared the observed p-values to those obtained from permuted data (as we did for our DEG analysis). The p-values of the permuted data resemble the null expectation, confirming that the differential peaks between high and low AI are likely true biological differences rather than artifacts (e.g. batch effects) (Fig. S20, Supplementary Data 10). In total we identified >20,000 peaks across cell types with significant differential accessibility between the high and low AI groups (FDR<10%), however, as with gene expression, most differences were small (log_2_FC < 0.1) (Fig. S21). This indicates that differences in addiction-like behaviors between rats are associated with modest regulatory changes at a large number of sites.

The differential peaks can be subdivided into those where accessibility is higher (upregulated) or lower (downregulated) in the high AI rats (Fig. S21). In astrocytes, there were roughly equal numbers of up- and downregulated peaks, but the other cell types showed profound directional biases. Excitatory neurons were the most biased with only two detected downregulated peaks, and over 8000 upregulated peaks in the high AI group. Inhibitory neurons showed the opposite bias with over 4000 downregulated peaks but only ∼500 upregulated peaks in the high AI group (Fig. S21). These biases likely reflect differences in the activity of specific TFs that control large transcriptional programs.

To determine whether the differential chromatin accessibility is consistent with the differential gene expression, we tested whether the promoters of DEGs are enriched for differential accessibility. We overlapped the significant differentially accessible chromatin peaks in each cell type with the promoters of significant DEGs and computed a log odds ratio (log_2_OR) as a measure of enrichment. Across all of the major cell types, differentially accessible peaks are enriched (FET, p<0.05) at the promoters of DEGs compared to non-DEGs (Fig. 5b, Table S4). We also examined chromatin accessibility at promoter regions for genes belonging to the oxidative phosphorylation pathway because genes within this pathway were enriched for gene expression differences between high vs. low AI rats in a majority of cell types. These genes are also significantly enriched for differentially accessible promoter peaks in inhibitory neurons, excitatory neurons and oligodendrocytes (Fig. S22, Table S5). These findings confirm that the observed differences in chromatin accessibility and gene expression are concordant.

The genomic annotations of the significant differential peaks showed that 3.19% of these regions were annotated as promoter or TSS regions (Fig. S23). While this is a small percentage of the peaks, it is consistent with other studies^26^. We then studied the subset of significant differential peaks in each cell type by examining their genomic annotations to determine if they were enriched for promoter/TSS regions compared to the set of all peaks. We observed that differentially accessible peaks were highly enriched in promoter regions, occurring at least four times more frequently than expected given the genomic annotations of all accessible chromatin regions in most of the major cell types (FET, FDR<10%) (Fig. 5c, Table S6). This enrichment may indicate that long-term changes in chromatin associated with addiction-like behaviors are more concentrated at promoters, or that we have greater statistical power to detect changes at promoters, due to larger effect sizes or greater overall chromatin accessibility.

We hypothesized that differences in chromatin accessibility between high and low AI rats are caused by differential TF activity. To test this hypothesis, we analyzed the snATAC-seq data using ChromVar (Supplementary Dataset 11), which identifies TF motifs associated with differential accessibility using sparse single cell data^90^. A large number of motifs have significant differences in accessibility between the high and low AI rats, and since many TFs recognize similar motifs, we grouped them into motif clusters and summarized results across cell types (Fig. 5d).

The motif cluster with the most significant difference in accessibility between high and low AI rats contains motifs for basic helix-loop-helix (bHLH) TFs. This motif cluster has substantially higher accessibility within the excitatory neurons of high AI rats compared to low AI rats (deviance 3.8, p=1e-280), as well as a modest increase in accessibility in inhibitory neurons (deviance 0.38, p=1e-34) (Fig. 5e-g). The top-ranked motifs in this cluster all harbor the sequence CAGATGG, which is a close match to binding site motifs for multiple neuronal pioneer TFs including those of the bHLH, RFX and FOX families^91,92^. Thus, the widespread increases in chromatin accessibility in excitatory neurons of high AI rats could reflect increased activity of pioneer TFs that recruit chromatin remodelers. However, we did not observe corresponding upregulation in the expression of genes encoding the TFs belonging to these clusters (Supplementary Data 5, Supplementary Data 11), suggesting that a different mechanism might affect their activity.

We noticed that many motif clusters with increased accessibility in the neurons of high AI rats have decreased accessibility in oligodendrocytes (Fig. 5d-g). Prominent among these motif clusters are those containing FOX and RFX motifs (Fig. 5d-g).

Several motif clusters also have opposite effects between excitatory and inhibitory neurons. SOX motifs have decreased accessibility in high AI rats in excitatory neurons but increased accessibility in all other major cell types including inhibitory neurons (Fig. 5d). MEF2 and Fos (AP1) motifs all have increased accessibility in the excitatory neurons of high AI rats but decreased accessibility in inhibitory neurons (Fig. 5d). AP1 and MEF2 motifs are of particular interest because they are associated with addiction^93–96^ and their expression changes in the brain following chronic exposure to cocaine and other drugs^97–101^. Consistent with these results, we observed that the expression of TFs of the AP1, including Fosl1, Fos, Jun, Junb, and Jund, was decreased in high versus low AI rats (Fig. S24), suggesting that differences in their expression level affect their transcriptional activity. While our analysis cannot pinpoint the precise TFs involved, it implicates many motif clusters that are associated with addiction-like behaviors across thousands of regulatory regions and in a cell type-specific manner.

Accessible chromatin regions harbor cell type-specific regulatory elements^102,103^, and enrichment analyses that measure intersections between ATAC-seq peaks and GWAS signals can yield insight into the mechanisms by which genetic variants confer risk^104^. However, cell type-specific measurements of chromatin accessibility are difficult to obtain from human brain tissues. To assess whether our rat snATAC-seq data is meaningful for interpreting human addiction-related traits, we mapped the accessible chromatin peaks to the human reference genome and performed cell type-specific LD score regression^105^. We chose to use summary statistics from well-powered GWAS for alcohol and tobacco use^106,107^ because there is significant genetic overlap among GWAS for all known substance use disorders^108^ and because available GWAS for cocaine use disorder are much smaller and less powerful. We found significant enrichments (FDR<10%) of SNP heritability in every trait tested in almost every cell type (Fig. 5h), with the most significant enrichments in neurons, astrocytes, oligodendrocytes and OPCs. Overall, these results support the hypothesis that, despite the millions of years of evolution separating humans and rats, the regulatory architecture of HS rats is relevant for human addiction-related traits.

## Discussion

To better understand the molecular basis of addiction and illuminate long-term changes in gene regulation and chromatin accessibility associated with chronic drug use, we have generated an atlas of single-cell gene expression and chromatin accessibility in the amygdala of rats that showed divergent cocaine addiction-like behaviors after a prolonged period of abstinence. Our dataset is the largest resource of cell types in the mammalian amygdala, with over 163,000 nuclei in our snRNA-seq dataset and 81,000 nuclei in our snATAC-seq dataset (Fig. 2a-b). The snATAC-seq dataset provides the first map of cell type-specific regulatory elements in the amygdala, which has allowed us to identify TF motifs that may drive addiction-related processes.

Previous single cell transcriptomic studies have focused on the effects of acute passive treatment with psychoactive drugs in rodents^37,38^, which cannot fully capture the motivational processes underlying addiction. In contrast, our behavioral protocol involves extended access to cocaine IVSA and reflects several key aspects of cocaine addiction, including escalation of drug use, enhanced motivation for drug seeking and taking, and persistent drug use despite adverse consequences, which might reflect compulsive-like drug consumption^109^. In addition, using an outbred population of rats with divergent addiction-like traits allowed us to correlate molecular differences not only a high AI phenotype, which reflects vulnerability to severe addiction-like phenotypes, but also to a low AI phenotype, which reflects resiliency to developing behaviors that are hallmarks of addiction. Thus, our study is the first to examine long-term molecular changes in distinct brain cell populations following abstinence from chronic voluntary cocaine use in vulnerable and resilient rats.

One striking finding from our study is that there were thousands of significant cell type-specific differences in gene expression and chromatin accessibility between high and low AI rats, with strong biases in the direction of regulation of open chromatin regions in several major cell types (Fig. S16, S17, S21). Most of these differences were small, which suggests that cocaine addiction-related behaviors may result from the combined action of many small effects on gene expression and chromatin accessibility. Because the HS rats are genetically diverse, the molecular differences between high and low AI rats could arise from genetic differences or from the consumption of different quantities of cocaine. These results are consistent with a polygenic model wherein addiction-like behaviors would result from the collective action of a large number of genetic risk loci with small individual effects. This is a plausible explanation because of the high genetic diversity in the HS rats and because complex traits in humans are highly polygenic^105,110^. Further support for this hypothesis comes from the observation that the majority of DEGs have eQTLs identified in HS rat brains^70^ (Fig. S17), including DEGs such as *Kcnq3, Fkbp5* and *Sgk1* (Fig. 3a-e).

Alternatively, the effects could be mediated by a relatively small number of TFs that affect many downstream genes and chromatin sites. Because some of the motifs with the strongest chromatin accessibility differences (Fig. 5e-h) are recognized by pioneer TFs (e.g. BHLH, Sox, Fox), it is tempting to speculate that widespread differences in accessibility are due to pioneer TFs, which have an intrinsic ability to modify chromatin^111^. These explanations are not mutually exclusive and it is likely that some differences are caused by eQTLs while others are caused by differences in the activity of upstream regulators (which themselves may be affected by genetics or other factors).

In an effort to uncouple pre-existing genetically controlled gene expression differences from cocaine-induced neuroadaptations, we performed an analysis comparing our observed DEGs to differences in expression obtained from genotype-based prediction models. This allowed us to leverage the genotype data from our cohort of genetically diverse HS rats. We found significant correlations in observed versus predicted differential gene expression between high vs. low AI rats, suggesting that genetics does play a role in long-term transcriptional neuroadaptations that are observed following cocaine use. While the correlation metrics we obtained from our analysis are modest, this is expected due to three limitations of the predictive model. First, the models are trained on whole brain tissue and do not have the same cell type-specific resolution as our snRNA-seq data. Second, the size of the cohort on which the predictive models were trained was quite modest. Third, the models are only capable of capturing a small fraction of variation in expression and do not account for other influences on gene expression. Finally, it is likely that the DEGs we discovered are biased towards highly expressed genes, and eQTLs are less detectable in genes with very low expression. While these limitations make it difficult to quantify how much of the variance in expression is due to genetics, it establishes that at least some of the differences are due genetic variation (Fig. S18). To properly uncouple pre-existing genetically controlled gene expression differences from cocaine-induced neuroadaptations would require a larger datasets of genotyped rats. One way this could be accomplished is through the use of polygenic risk scores for addiction-related traits, which will become possible as more rat behavioral GWAS are completed^43,45–47,112^.

Human and animal studies have provided genetic and behavioral evidence that alterations in GABAergic transmission are involved in addiction^2,113–117^. Consistent with prior findings showing that GABAergic transmission is enhanced following excessive cocaine use^118^, our differential gene expression analysis showed enrichment of genes belonging to the GABAergic synapse pathway (Fig. 3f) and our electrophysiology results provided evidence for enhanced GABAergic transmission in the high AI rats (Fig. 4b). Moreover, we found that inhibition of GLO1, the enzyme responsible for MG metabolism, normalizes electrophysiological (Fig. 4c-f) and behavioral differences (Fig. 4h) associated with severe addiction-like behaviors. Specifically, while pBBG normalized the increased GABA transmission in electrophysiological recordings for both low and high AI rats (Fig. 4c), it had a normalizing effect on the drug-seeking behaviors in high AI rats but not low AI rats (Fig. 4h). This suggests that the inhibitory effects of pBBG on relapse-like behaviors depend on a given threshold of GABAergic transmission. These results corroborate previous findings that MG acts as an endogenous competitive agonist for GABA_A_ receptors^113,119^. GABA_A_ receptor agonists used in the context of cocaine-seeking behavior have shown contrasting results leading to both reduction or increase in cocaine-seeking behaviors^120–128^. Since MG is generated in proportion to glycolytic activity of nearly every cell and does not accumulate in synaptic vesicles, it diffuses and may activate GABA_A_ receptors at synaptic and extra synaptic sites; thus, manipulating the endogenous levels of MG by GLO1 inhibition probes a mechanism of GABA_A_ receptor regulation that is fundamentally different from the canonical modulation of synaptic GABA_A_ receptors. In our study, electrophysiological recordings suggest that there is an increase in GABAergic transmission without changes in postsynaptic currents in the CeA; thus, we speculate that MG-based pharmacological manipulations may alter presynaptic GABA_A_ receptor function, reducing GABA release at inhibitory terminals and suppressing inhibitory connections within the CeA. Consistently, previous studies demonstrated that the activation of presynaptic GABA_B_ receptors suppresses inhibitory connection within the CeA^84^ and that negative regulation of GABAergic transmission can occur through a presynaptic mechanism^85^. An alternative scenario is that the magnitude of effects is not sufficient to cause detectable changes in amplitude. Overall, these results offer a promising pharmacological target for improving therapeutic approaches for cocaine addiction that was identified by our single cell analysis of the amygdala in high and low AI rats.

While the pharmacological inhibition experiments are not cell type-specific, the pathway enrichment analysis of the transcriptomic data suggest that GABAergic synapse-related genes may be specific to Cck+Vip+ and Nos1+ subtypes of inhibitory neurons. Previous studies manipulating GLO1 activity directly in the mouse amygdala by transgenic expression of *Glo1* or MG microinjection were sufficient to reduce anxiety-like behaviors^129^. Future experiments targeting specific subregions or cell types of the amygdala will be necessary to further characterize the effects of GLO1 inhibition on cocaine addiction-related phenotypes.

The results from the GLO1 inhibition experiments indicate a close connection between localized energy metabolism and neurotransmission^130^. Moreover, genes which are differentially regulated in high versus low AI rats are enriched in pathways related to energy metabolism, including glycolysis, pyruvate metabolism, and oxidative phosphorylation (Fig. 3f). Most notably, the expression levels of genes related to oxidative phosphorylation, which determines cellular ATP levels^131^, are altered across most amygdalar cell types. Not only is ATP crucial for sustaining electrophysiological activity and cell signaling in the brain^132,133^, but it is also required for ATP-dependent chromatin remodeling events initiated by pioneer TFs^134^. This could potentially explain why excitatory and inhibitory neurons show opposite directions of regulation in chromatin accessibility (Fig. S21) and in the enrichment of DEGs in the oxidative phosphorylation pathway (Fig. 3f). In combination, these observations suggest that an altered metabolic state within the amygdala impacts multiple cellular processes that are involved in vulnerability to and development of addiction. Future experiments that directly manipulate the expression of specific metabolic enzymes or pioneer TFs in a cell type-specific manner will be necessary to fully elucidate their role in addiction.

In conclusion, the cellular atlas created by this study is a valuable resource for understanding cell type-specific gene regulatory programs in the amygdala and their role in the development of cocaine addiction-related behaviors. Our results emphasize the contribution of cellular energetics and the GABA_A_-mediated signaling to the enduring effects of cocaine use, which led us to perform experiments that manipulate GABA_A_ transmission via the pharmacological inhibition of GLO1 and identify a novel potential target for treatment of cocaine addiction. We anticipate that future studies will utilize our data to further investigate novel cell type-specific mechanisms involved in addiction.

## Methods

### Experimental

#### Animals

All protocols were reviewed and approved by the institutional Animal Care and Use Committee at the University of California San Diego. HS rats (Rat Genome Database NMcwiWFsm #13673907, sometimes referred to as N/NIH) which were created to encompass as much genetic diversity as possible at the NIH in the 1980’s by outbreeding eight inbred rat strains (ACI/N, BN/SsN, BUF/N, F344/N, M520/N, MR/N, WKY/N and WN/N) were provided by Dr. Leah Solberg Woods (Wake Forest University School of Medicine). To minimize inbreeding and control genetic drift, the HS rat colony consists of 64 or more breeder pairs and is maintained using a randomized breeding strategy, with each breeder pair contributing one male and one female to subsequent generations. To keep track of the rats, their breeding, behavior, organs and genomic info, each rat received a chip with an RFID code. Rats were shipped at 3-4 weeks of age, kept in quarantine for 2 weeks and then housed two per cage on a 12 h/12 h reversed light/dark cycle in a temperature (20-22°C) and humidity (45-55%) controlled vivarium with ad libitum access to tap water and food pellets (PJ Noyes Company, Lancaster, NH, USA). We used 46 HS rats for the behavioral experiments presented in Fig. 1, of which 20 male rats (high and low AI) were used for the generation of snRNA-seq and snATAC-seq data, along with 11 naive male rats. Additionally, 26 of these 46 behaviorally phenotyped rats (13 female, 13 male) were used for the cue-induced reinstatement experiments. For snRNA-seq, we used 19 male rats (6 high AI, 6 low AI, 7 naive). For the snATAC-seq, we used 12 male rats (4 high AI, 4 low AI, 4 naive). In addition, we used a different cohort of 15 female and male rats (5 high AI, 5 low AI, 5 naive) for the electrophysiology experiments.

#### Drugs

Cocaine HCl (National Institute on Drug Abuse, Bethesda, MD, USA) was dissolved in 0.9% saline (Hospira, Lake Forest, IL, USA) and administered intravenously at a dose of 0.5 mg/kg/infusion as described below. pBBG was synthesized in the laboratory of Prof. Dionicio Siegel (University of California San Diego, Skaggs School of Pharmacy and Pharmaceutical Sciences). pBBG was dissolved in a vehicle of 8% dimethylsulfoxide, 18% Tween-80, and 74% distilled water and administered intraperitoneally 30 minutes before the test session.

#### Brain Samples

Brain tissues were obtained from the cocaine brain bank at UCSD^39^, which collects tissues from HS rats that are part of an ongoing study of addiction-like behavior^43^. We used 31 HS rats for generating the single-nucleus sequencing data reported in this study, which included 20 rats that had low or high AI for cocaine addiction-related behaviors, using behavioral methods previously described^48^ were kept in their home cages and never subjected to the catheter implantation or the behavioral protocol for cocaine IVSA. Brain tissues were collected after four weeks of abstinence from cocaine self-administration, which has been used in prior studies to examine long-lasting effects of self-administration^47,135–140^. Brain tissues were extracted and snap-frozen (at −30°C). Cryosections of approximately 500 microns (Bregma -1.80 - 3.30mm) were used to dissect the amygdala on a −20°C frozen stage, including the CeA, BLA, and medial amygdala from both hemispheres. Punches from three sections were combined for each rat. In addition, 6 ACI/EurMcw rats were used for the dissection of the CeA and BLA.

#### Single-cell library preparation, sequencing, and alignment

snRNA-seq libraries from the whole amygdala tissues were performed by the Center for Epigenomics, UC San Diego using the Droplet-based Chromium Single-Cell 3’ solution (10x Genomics, v3 chemistry). Briefly, frozen tissue was homogenized via glass dounce. Nuclei were then resuspended in 500 µL of nuclei permeabilization buffer (0.1% Triton-X-100 (Sigma-Aldrich, T8787), 1X protease inhibitor, 1 mM DTT, and 1U/µL RNase inhibitor (Promega, N211B), 2% BSA (Sigma-Aldrich, SRE0036) in PBS). Sample was incubated on a rotator for 5 min at 4°C and then centrifuged at 500 rcf for 5 min (4°C, run speed 3/3). Supernatant was removed and pellet was resuspended in 400 µL of sort buffer (1 mM EDTA 0.2 U/µL RNase inhibitor (Promega, N211B), 2% BSA (Sigma-Aldrich, SRE0036) in PBS) and stained with DRAQ7 (1:100; Cell Signaling, 7406). Up to 75,000 nuclei were sorted using a SH800 sorter (Sony) into 50 µL of collection buffer consisting of 1 U/µL RNase inhibitor in 5% BSA; the FACS gating strategy sorted based on particle size and DRAQ7 fluorescence. Sorted nuclei were then centrifuged at 1000 rcf for 15 min (4°C, run speed 3/3) and supernatant was removed. Nuclei were resuspended in 35 µL of reaction buffer (0.2 U/µL RNase inhibitor (Promega, N211B), 2% BSA (Sigma-Aldrich, SRE0036) in PBS) and counted on a hemocytometer. 12,000 nuclei were loaded onto a Chromium Controller (10x Genomics). Libraries were generated using the Chromium Single-Cell 3′ Library Construction Kit v3 (10x Genomics, 1000075) with the Chromium Single-Cell B Chip Kit (10x Genomics, 1000153) and the Chromium i7 Multiplex Kit for sample indexing (10x Genomics, 120262) according to manufacturer specifications. cDNA was amplified for 12 PCR cycles.

For snATAC-seq libraries from the whole amygdala tissues, nuclei were purified from frozen tissues using an established method^141^. Briefly, frozen amygdala tissue was homogenized using a 2 ml glass dounce with 1 ml cold 1x Homogenization Buffer (HB). The cell suspension was filtered using a 70 μm Flowmi strainer (BAH136800070, Millipore Sigma) and centrifuged at 350g for 5 min at 4°C. Nuclei were isolated by iodixanol (D1556, Millipore Sigma) density gradient. The nuclei iodixanol solution (25%) was layered on top of 40% and 30% iodixanol solutions. Samples were centrifuged in a swinging bucket centrifuge at 3,000g for 20 min at 4°C. Nuclei were isolated from the 30-40% interface. Libraries were generated using the Chromium Next GEM Single Cell ATAC v1.1 (10x Genomics, PN-1000175) with the Chromium Next GEM Chip H Single Cell Kit (10x Genomics, 1000162) and the Chromium i7 Multiplex Kit for sample indexing (10x Genomics, 1000212) according to manufacturer specifications. DNA was amplified for 8 cycles.

For snRNA libraries from BLA and CeA, frozen brain tissues were obtained from the ACI/EurMcw rat strain, one of the HS rat founder strains. For nuclei isolation, brain punches from 3 rats for each region were pooled and homogenized in homogenization buffer (0.26 M sucrose, 0.03 M KCl, 0.01 M MgCl2, 0.02 M Tricine-KOH pH 7.8, 0.001 M DTT, 0.5 mM spermidine, 0.15 mM Spermine, 0.3% NP40) using with 1ml glass Dounce homogenizers. The homogenate was filtered with a 70-um strainer filter (SP Bel-Art, cat no 136800070) and centrifuged for 5 minutes at 350 RCF. The nuclei were purified with an iodixanol gradient (Sigma-Aldrich # 92339-11-2) by layering a 25% Iodixanol-nuclei mixture on top of 30% and 40% Iodixanol solutions. After centrifugation at 4°C 3,000 RCF for 20 minutes, nuclei were collected from the 30-40% interface. Nuclei were washed in ATAC-RSB-Tween buffer (0.01 M Tris-HCl pH 7.5, 0.01 M NaCl, 0.003 M MgCl2, 0.1% Tween-20) and then resuspended in 1X nuclei buffer (10X genomics, PN 2000207). 12,000 nuclei were loaded on the 10X Genomics Chromium Controller for GEM generation. RNAse inhibitors (Roche Diagnostics, # 03335402001) were added to all buffers (1U/ul). snRNA-seq library was performed using the Chromium Next GEM Single Cell Multiome Reagent Kit A (# 1000282) following Chromium Next GEM Single Cell Multiome ATAC + Gene Expression Reagent Kits User Guide (10X Genomics). After the transposition reaction, nuclei were encapsulated and barcoded. Next-generation sequencing libraries were constructed following the User Guide.

For each library type, SPRISelect reagent (Beckman Coulter, B23319) was used for size selection and clean-up steps. Final library concentration was assessed by Qubit dsDNA HS Assay Kit (Thermo-Fisher Scientific) and post library QC was performed using Tapestation High Sensitivity D1000 (Agilent) to ensure that fragment sizes were distributed as expected. Final libraries were sequenced using the NovaSeq6000 (Illumina).

#### Behavioral experiments

Intravenous catheterization and behavioral testing of rats used for the generation of snRNA-seq and snATAC-seq were carried out using an established protocol of extended access to cocaine IVSA, progressive ratio (PR) testing, and foot shock, as reported previously^39,48,49^. Briefly, after surgical implantation of intravenous catheters and a week of recovery, HS rats were trained to self-administer cocaine (0.5 mg/kg/infusion) in 10 short access (ShA) sessions (2h/day, 5 days per week). Next, the animals were allowed to self-administer cocaine in 14 long access (LgA) sessions (6h/day, 5 days/week) to measure the escalation of drug intake (Fig. 1e). Following the escalation phase, rats were screened for motivation using the PR test and for persistent drug-seeking despite adverse consequences using contingent foot-shock. The breakpoint (Fig. 1f) was defined as the maximal number of presses completed before a 60-minute period during which a rat does not complete the next schedule. Rats were classified as vulnerable (high AI), defined by having high addiction-like behavior, versus resilient (low AI), defined as having low addiction-like behavior, via a median split^51,52^. AI was computed by averaging normalized measurements (z-scores) for the three behavioral tests: escalation of drug intake, motivation, and compulsive-like behavior, or drug taking despite adverse consequences^142^ (Fig. 1c-d). The z-scores were calculated as Z = (x-μ)/σ, where x is the raw value, μ is the mean of the cohort, and σ is the standard deviation of the cohort. Brain tissues were collected after four weeks of abstinence. For the pBBG studies, we used rats with low and high AI distinct from those used for the snRNA-seq and snATAC-seq experiments. Four weeks after the last drug self-administration session, the rats were placed back in the self-administration chambers without the availability of cocaine. The number of responses to the previous drug-paired lever (cocaine seeking behavior) was measured 30 minutes after intraperitoneal injection of pBBG (15 mg/kg/ml) or its vehicle, in a Latin square design. The 30 minutes time point was selected based on a previous study^86^. Data were analyzed using Prism 9.0 software (GraphPad, San Diego, CA, USA). Self-administration data were analyzed using repeated-measures analysis of variance (ANOVA) or mixed effect model followed by Bonferroni post-hoc tests when appropriate. For pairwise comparisons, data were analyzed using the unpaired *t*-test. The data are expressed as mean ± SEM unless otherwise specified. Values of p < 0.05 were considered statistically significant.

#### Electrophysiology

Slices of the CeA were prepared from rats after 4 weeks of protracted abstinence from cocaine IVSA following the same behavioral protocol described above or age-matched naive rats. High AI (n=5), low AI (n=5) and naive (n=5) rats were used for patch clamp baseline recordings. These rats were distinct from those used for snRNA-seq and snATAC-seq. Slices from each group were also used to record iPSCs after pBBG treatment. The naive rats received sham IV surgery. The rats were deeply anesthetized with isoflurane and brains were rapidly removed and placed in oxygenated (95% O_2_, 5% CO_2_) ice-cold cutting solution that contained 206 mM sucrose, 2.5 mM KCl, 1.2 mM NaH_2_PO_4_, 7 mM MgCl_2_, 0.5 mM CaCl_2_, 26 mM NaHCO_3_, 5 mM glucose, and 5 mM Hepes. Transverse slices (300 μm thick) were cut on a Vibratome (Leica VT1200S; Leica Microsystems) and transferred to oxygenated artificial cerebrospinal fluid (aCSF) that contained 130 mM NaCl, 2.5 mM KCl, 1.2 mM NaH_2_PO_4_, 2.0 mM MgSO_4_·7H_2_O, 2.0 mM CaCl_2_, 26 mM NaHCO_3_, and 10 mM glucose. The slices were first incubated for 30 min at 35°C and then kept at room temperature for the remainder of the experiment. Individual slices containing CeA were transferred to a recording chamber that was mounted on the stage of an upright microscope (Olympus BX50WI). The slices were continuously perfused with oxygenated aCSF at a rate of 3 mL/min. Neurons were visualized with a 60X water-immersion objective (Olympus), infrared differential interference contrast optics, and a charge coupled device camera (EXi Blue; QImaging). Whole-cell recordings were performed using a Multiclamp 700B amplifier (10 kHz sampling rate, 10 kHz low-pass filter) and Digidata 1440A and pClamp 10 software (Molecular Devices). Patch pipettes (4–6 MΩ) were pulled from borosilicate glass (Warner Instruments) and filled with 70 mM KMeSO_4_, 55 mM KCl, 10 mM NaCl, 2 mM MgCl_2_, 10 mM Hepes, 2 mM Na-ATP, and 0.2 mM Na-GTP. Pharmacologically isolated sIPSCs were recorded in the presence of the glutamate receptor blockers, CNQX (Tocris #0190) and APV (Tocris #189), and the GABA-B receptor antagonist CGP55845 (Tocris #1246). Experiments with a series resistance of >15 MΩ or >20% change in series resistance were excluded from the final dataset. pBBG (2.5uM) was acutely applied in the bath. The frequency, amplitude, and kinetics of sIPSCs were analyzed using semi automated threshold-based mini detection software (Easy Electrophysiology) and visually confirmed. Data were analyzed using Prism 9.0 software (GraphPad, San Diego, CA, USA) with one-way ANOVA followed by post hoc Tukey HSD test or with paired t-tests. The data are expressed as mean ± SEM unless otherwise specified. Values of p < 0.05 were considered statistically significant.

### Computational

#### Alignment of snRNA-seq and snATAC-seq reads

Raw base call (BCL) files were used to generate FASTQ files using Cell Ranger 3.1.0 for snRNA-seq data, Cellranger ATAC v.2.0.0 for snATAC-seq data, and Cell Ranger ARC v.2.0.0 for processing Chromium Single Cell Multiome ATAC + Gene Expression sequencing data. For demultiplexing raw base call (BCL) files generated by the sequencers into FASTQ files, we used the ‘cellranger mkfastq’ command for RNA-seq reads, ‘cellranger-atac mkfastq’ for ATAC-seq reads, and ‘cellranger-arc mkfastq’ for the CeA and BLA samples which were generated using the multiome kit^143,144^. Next, we used ‘cellranger count’ and ‘cellranger-atac count’ to align the snRNA-seq and snATAC-seq reads, respectively, to a custom rat reference genome based on the rn6 reference genome downloaded from the UCSC genome browser^145–147^. The reference genome files for cell ranger were created from FASTA and genome annotation files for Rattus norvegicus Rnor_6.0 (Ensembl release 98)^148^ and JASPAR2022 motifs^149^. BLA and CeA samples were aligned to the same reference genome using ‘cellranger-arc count’. We then filtered cells based on quality control metrics and performed barcode and UMI counting for the RNA-seq and ATAC-seq reads.

#### Quality control and preprocessing of snRNA-seq data

All snRNA-seq preprocessing was performed with Seurat v3.2.3^56^. For each sample, we loaded the filtered feature barcode matrices containing only detected cellular barcodes returned by ‘cellranger count’ into Seurat. We computed the number of unique genes detected in each cell (nFeature_RNA); the total number of molecules detected within a cell (nCount_RNA); and the percentage of reads mapping to the mitochondrial genome (percent.mt) (Fig. S1-3, Supplementary Data 1, Supplementary Data 3). nFeature_RNA is informative because low-quality cells or empty droplets will typically have very low numbers of detected genes while doublets or multiplets will exhibit very high gene counts. nCount_RNA is a metric that correlates with nFeature_RNA. We examined percent.mt because low-quality or dying cells typically exhibit a high degree of mitochondrial contamination. We removed all cells for which the value of any of these metrics was more than three standard deviations from the mean within the sample (*x* > μ ± 3σ). Next, we normalized the count data for each sample using sctransform^150^ with percent.mt as a covariate.

#### Integrating snRNA-seq data across samples and clustering

To integrate snRNA-seq data across all of our samples, we used reciprocal principal component analysis (RPCA), as implemented in Seurat^56,151^. First, we identified 2000 highly variable features (genes) across all of the samples to use as integration features using the ‘SelectIntegrationFeatures()’ function, which we passed as anchor features (‘anchor.features’) to the ‘PrepSCTIntegration()’ function. Next, we performed dimensionality reduction with PCA on each sample using ‘RunPCA()’. After this, we ran the ‘FindIntegrationAnchors()’ function to find a set of anchors between the list of Seurat objects from all of our samples using the same anchor features passed to ‘PrepSCTIntegration()’, specifying RPCA as the dimensional reduction method to be performed for finding anchors (‘reduction = rpcà) and the first 30 RPCA dimensions to be used for specifying the k-nearest neighbor search space. Two rats (1 high AI, 1 low AI) were used as reference samples for the integration. We used the resulting anchor set to perform dataset integration across all of our samples using ‘IntegrateData()’. We clustered the integrated dataset by constructing a K-nearest neighbor (KNN) graph using the first 30 PCs followed by the Louvain algorithm, implemented in Seurat using the ‘FindNeighbors()’ function followed by ‘FindClusters()’. Finally, we ran PCA once again on the integrated dataset and visualized the data using uniform manifold approximation and projection (UMAP). Visualization of the integrated data in two-dimensional space indicated that batch correction was successful (Fig. S9a-c). To compare CeA and BLA subregion samples with the whole amygdala, we subsampled whole amygdala samples from the naïve rats in our study and performed the same integration technique. The integrated subregion data was visualized using UMAP.

#### Cell type assignment for snRNA-seq data

We identified marker genes of each cluster in our integrated snRNA-seq dataset using MAST^152^, implemented with the ‘FindMarkers()’ function in Seurat. Cell type identities were assigned based on expression of known marker genes of cell types known to be found in the amygdala.

#### Cell type-specific gene expression analysis for snRNA-seq data

Within each cell type, we tested for DEGs between the high AI rats and the low AI rats. We used the negative binomial test^88,89^ implemented with the ‘FindMarkers()’ function in Seurat to identify differential expression between groups, using percent.mt and the library prep date as covariates. We used the avg_log2FC value returned by the ‘FindMarkers()’ function as an estimate of effect size. We did not pre-filter genes for testing based on log-fold change or minimum fraction of cells in which a gene was detected. This approach allowed us to detect weaker signals because we tested all observed genes in the dataset. Multiple testing correction was performed using the Benjamini-Hochberg method and we used a false discovery rate of 10% as a significance threshold (FDR<10%). Permutation tests were performed using the same methods, covariates, and filtering options but with shuffled addiction index labels. Results from permuted and unpermuted data were compared by visualizing the distributions of uncorrected p-values with QQ-plots (Fig. S15, Supplementary Data 6).

We used ClusterProfiler^153^ to perform gene set enrichment analysis (GSEA) of KEGG pathways for DEGs from each cell type. A ranked list of the avg_logFC values for all genes evaluated with our negative binomial test was given as input to GSEA. Multiple testing correction for GSEA results was performed using Benjamini-Hochberg adjustment, with statistical significance assessed at a FDR<10%.

All rat eQTLs described in the paper come from the RatGTEx portal (https://ratgtex.org/download/). Specifically, we downloaded their table of conditionally independent cis-eQTLs, which only includes eQTLs passing a significance threshold of FDR<0.05. We examined cis-eQTLs in the following brain tissues: BLA, Brain, IL, LHb, NAcc, NAcc2, OFC, PL, PL2. For each cis-eQTL in the database, only the top associated SNP is given, but some genes have more than one cis-eQTL in a tissue, meaning there are multiple loci with statistically independent associations with the gene’s expression. We measured enrichment of significant DEGs (FDR<10%) that also had eQTLS in the rat brain using the Chi-squared test, implemented using the ‘chisq.test()’ function in R.

To obtain bootstrap distributions of DEG effect sizes, we resampled nuclei with replacement 1000 times. Resampling was performed separately for nuclei from high and low AI rats so that the sample size of each set remained consistent. For each bootstrap iteration we recorded the p-value and the coefficients for the hi/low AI condition from the negative binomial regression performed by Seurat’s ‘FindMarkers()’ function. We then rescaled the coefficient to be in units of log2 fold change. We note that the log2FC estimates obtained by this method correspond to the p-values, but differ slightly from Seurat’s avg_log2FC estimates because Seurat’s avg_log2FC calculations introduce a pseudocount and do not consider the effects of covariates. The distribution of resulting bootstrap fold-change estimates and q-values were visualized as violin plots using Plotly in Python (Fig. 3c-e).

#### Comparing observed gene expression differences to predicted gene expression differences based on cis-genetic variation

To estimate the genetic component of gene expression variation in the brain, conditionally independent cis-eQTLs and their allelic fold change (aFC) estimates for whole brain hemisphere tissue were downloaded from the RatGTEx Portal (https://ratgtex.org/download/). Using aFC as effect size, gene expression was predicted from genotypes using eQTL linear models^77^ (https://github.com/PejLab/gene_expr_pred). Predicted relative expression was thus obtained for 26 rats whose genotypes were available, and only for genes with at least one significant cis-eQTL. Genes with zero-variance predictions were removed, resulting in predictions for 8,997 genes. To further prioritize genes by estimated prediction accuracy, gene expression was predicted for the same 339 rats that were used to map the whole brain hemisphere eQTLs. Pearson correlation *r*^2^ was calculated between those predictions and observed log-TPM expression from the same rats. We then measured the difference between mean predicted expression in high vs low AI rats and compared it against the avg_logFC estimates obtained by Seurat’s FindMarkers() function. Spearman’s correlation coefficient (*ρ*) was calculated between the difference in mean predicted expression and the observed avg_logFC. We performed these tests multiple times using different *r*^2^ cutoffs for the gene expression prediction models to filter genes (Table. S3). Spearman’s correlation coefficients (*ρ*) and the associated p-values were calculated using ‘scipy.stats.spearmanr()’. Confidence intervals were calculated using the formula 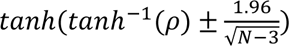. Spearman *ρ* confidence intervals were visualized using Plotly in Python.

#### Quality control and preprocessing of snATAC-seq data

All snATAC-seq data preprocessing was performed with MACS2^87^ (for peak calling) and Signac^57^. Although peak calling is performed during alignment by ‘cellranger-atac count’, we chose to call peaks separately using MACS2 because Cell Ranger’s peak calling function has been observed to merge multiple distinct peaks into a single region^154^. To minimize loss of informative features for clustering and downstream analysis, we first called peaks on the snATAC-seq BAM files for each rat with MACS2 (‘macs2 callpeak -t {input} -f BAM -n {sample} --outdir {output} {params} --nomodel --shift -100 --ext 200 --qval 5e-2 -B --SPMR’). We confirmed that MACS2 calls more peaks and peaks with smaller widths compared to Cell Ranger (Fig. S25). Next, we merged overlapping peaks across all of our samples to generate a combined peak set using BEDtools^155^ (‘bedtools mergè). We generated a new peak by barcode matrix for each sample using this combined peak set and all detected cellular barcodes using the ‘FeatureMatrix()’ function in Signac. We used these new matrices to create ChromatinAssay objects in Signac with the BSgenome.Rnorvegicus.UCSC.rn6^146^ reference genome using the ‘CreateChromatinAssay()’ function. From these ChromatinAssay objects we created Seurat objects with ‘CreateSeuratObject()’, which are compatible with functions from the Seurat package. We computed several quality control metrics for each sample: nucleosome banding pattern (nucleosome_signal); transcriptional start site (TSS) enrichment score (TSS.enrichment); total number of fragments in peaks (peak_region_fragments); and fraction of fragments in peaks (pct_reads_in_peaks) (Fig. S4-6, Supplementary Data 2, Supplementary Data 4). We removed all cells for which the value of any of these metrics was more than two standard deviations from the mean within the sample (*x* > μ ± 2σ). We removed one rat (FTL_463_M757_933000320046135) from our dataset, due to the very low number of detected fragments per cell in this sample (Fig. S26).

#### Integrating snATAC-seq data across samples and clustering

Each sample was normalized using term frequency-inverse document frequency (TF-IDF) followed by singular value decomposition (SVD) on the TF-IDF matrix using all features (peaks)^57,154^. The combined steps of TF-IDF followed by SVD are known as latent semantic indexing (LSI)^156,157^. Non-linear dimensionality reduction and clustering were performed using UMAP and KNN following the same procedure used, respectively, just as we did for the snRNA-seq data. We merged the data across all samples within Signac and repeated the process of LSI in the same manner as it was applied to individual samples. We then integrated the merged dataset using Harmony^158^ implemented by Signac, integrating over the sample library variable to account for the effects of different sequencing libraries used for different samples. We observed successful reduction of batch effects following integration Fig. S9d-f. We once again performed non-linear dimensionality reduction and clustering with UMAP and KNN, respectively. Notably, LSI, UMAP and KNN are used for visualization purposes; raw counts were used for downstream differential accessibility analyses.

#### Label transfer and cell type assignment for snATAC-seq data

We created a gene activity matrix for the integrated snATAC-seq data using the ‘GeneActivity()’ function in Signac. This counts the number of fragments per cell overlapping the promoter region of a given gene and uses that value as a gene activity score. Gene activity serves as a proxy for gene expression as gene expression is often correlated with promoter accessibility. The gene activity scores were log normalized using the ‘NormalizeData()’ function in Seurat with the normalization method set to ‘LogNormalizè and the scaling factor set to the median value of nCount_RNA across all cells, based on the gene activity scores. Cell type identities were assigned by integrating the snATAC-seq data with the integrated snRNA-seq data and performing label transfer^56^ as described in Signac. Briefly, this approach identifies shared correlation patterns in the gene activity matrix and the scRNA-seq dataset to identify matched biological states across the two modalities. The process returns a classification score for each cell for each cell type defined in the scRNA-seq data. Each cell was assigned the cell type identity with the highest prediction score. Additionally, by identifying matched cells in the snRNA-seq dataset, we were able to impute RNA expression values for each of the cells in our snATAC-seq dataset. This enabled us to perform correlative analyses of chromatin accessibility and gene expression in later downstream analyses, as it produced a pseudo-multimodal dataset.

#### Differential chromatin accessibility analysis of snATAC-seq data

Similar to our differential analyses of the snRNA-seq data, we tested for differentially accessible genomic regions between nuclei from the high versus low AI rats within each cell type. We used the negative binomial test^150,159^ implemented with the ‘FindMarkers()’ function from Seurat to model the raw snATAC-seq count data using peak_region_fragments, library batch date, and rat sample ID as covariates. Multiple testing correction was performed using Benjamini-Hochberg adjustment and a false discovery rate below 10% (FDR<10%) was used to determine statistical significance. Permutation tests were performed in the same manner as for the differential gene expression analyses (using the same statistical test and covariates with shuffled addiction index labels).

#### Partitioned heritability analysis

We downloaded summary statistics for the Liu et al. 2019 GWAS of tobacco and alcohol use^106^ and used the munge_sumstats.py script from LD Score (LDSC)^105^ to parse the summary statistics file into the proper format for downstream analyses. We used the sets of significant differential peaks (FDR<10%) for each cell type as foreground peaks and DNaseI hypersensitivity profiles for 53 epigenomes from ENCODE Honeybadger2. We used the UCSC liftOver tool to convert the foreground peaks from rn6 to hg19. There was no need to lift over the background peaks as Honeybadger2 is already in hg19. Next, we generated partitioned LD scores for the background and foreground regions. We used the make_annot.py script to make annotation files and the ldsc.py script to compute annotation-specific LD scores. We used the European 1000 Genomes Phase 3 PLINK^160^ files to compute the LD scores. Finally, using the baseline model and standard regression weights from the LDSC Partitioning Heritability tutorial, we ran a cell type-specific partitioned heritability analysis with the LD scores we computed.

#### Annotation of accessible chromatin regions

Before performing any differential analyses, we first used the annotatePeaks.pl script from the HOMER suite to annotate accessible chromatin regions and significant differential peaks (FDR<10%) for each cell type in our integrated dataset^161^. For each cell type, we performed a Fisher’s Exact Test to measure the enrichment of genomic regions annotated as a promoter region within the differential peaks compared to the set of all peaks in the dataset and observed significant results for all cell types tested. Specifically, we compared the ratio of peaks annotated as promoter regions to non-promoter regions in the significant differential peaks (FDR<10%) versus all other peaks.

#### Fisher’s Exact Tests

We first performed a Fisher’s Exact Test to measure enrichment of DEGs (FDR<10%) with differentially accessible promoters. We defined the latter as the case where the promoter region of a gene overlaps a significant differentially accessible peak (FDR<10%). We obtained gene coordinates from the TxDb.Rnorvegicus.UCSC.rn6.refGene annotation package and defined promoter regions as being 1500 bases upstream and 500 bases downstream of the TSS (‘promoters(genes(TxDb.Rnorvegicus.UCSC.rn6.refGene), upstream = 1500, downstream = 500)’). We then generated a confusion matrix from the following four values: the number of DEGs with differentially accessible promoters; the number of DEGs with non-differentially accessible promoters; the number of non-DEGs with differentially accessible promoters; and the number of non-DEGs with non-differentially accessible promoters. We then performed a Fisher’s Exact Test to measure enrichment of differentially accessible peaks (FDR<10%) which were annotated as TSS/promoter regions by HOMER (annotatePeaks.pl). We generated a confusion matrix from the following four values: the number of differential peaks with a TSS/promoter annotation; the number of differential peaks without a TSS/promoter annotation; the number of non-differential peaks (FDR>10%) with a TSS/promoter annotation; and the number of non-differential peaks (FDR>10%) without a TSS/promoter annotation.

#### Measuring differential activity of transcription factors with chromVAR

We measured cell type specific motif activities using chromVAR to test for per motif deviations in accessibility between nuclei from high versus low AI rats. Motif data was pulled from the JASPAR2020 database, and integrated with snATAC-seq data using the ‘AddMotifs()’ function in Signac. After adding motifs to our snATAC-seq dataset, we ran chromVAR with the ‘RunChromVAR()’ wrapper in Signac. Differential analysis of chromVAR deviation scores was performed using the Wilcoxon rank-sum test between high AI rats versus lowly addicted rats within each cell type. Differential testing was performed using Seurat’s ‘FindMarkers()’ function with the mean function set as ‘rowMeans()’ to calculate average difference in deviation scores between groups. Multiple testing correction was performed using Benjamini-Hochberg adjustment and a false discovery rate below 10% (FDR<10%) was used to determine statistical significance. Motif clusters were defined using the provided cluster numbers from JASPAR’s matrix clustering-results and the names of the clusters were annotated by hand based on which TFs were present in each cluster. When selecting clusters to visualize, we retrieved the top 50 motifs (FDR<10%) per cell-type and highlighted their respective clusters. Volcano plots and heatmap data were generated using Plotly in Python. Hierarchical ordering of heatmap clusters was generated with Plotly’s ‘figure_factory.create_dendrogram()’ function, which wraps the ‘cluster.hierarchy.dendrogram()’ function in SciPy.

## Data availability

The datasets generated in the current study are available through the Gene Expression Omnibus (GSE212417, token gpczagqerncrfmz).

The following publicly available datasets were used: Rattus norvegicus Ensembl v98 reference genome and genome assembly (Rnor_6.0); JASPAR2022 transcription factor binding profiles for vertebrates; ENCODE Honeybadger 2 ChIP-seq; 1000 Genomes European reference panel; Liu et al. 2019^106^ GWAS for tobacco and nicotine addiction; RatGTEx Portal tissue-specific cis-eQTLs.

The behavioral data of HS rats are available at https://cgord.org/

## Code availability

All code used to perform analyses in this paper can be found at https://github.com/mcvickerlab/sn_cocaine_rats.

## Supporting information

Supplementary Data Guide

## Acknowledgements

We thank S. Preissl from the UC San Diego School of Medicine Center for Epigenomics for technical assistance with the snRNA-seq library preparation. We thank J. Hightower for assistance with figures preparation. We thank P. Montilla-Perez, L. Maturin and P. Schweitzer for technical assistance with sample collection and equipment maintenance. We thank Leah C. Solberg Woods for HS rats breeding colony management. This work was supported by the National Institutes of Health (NIH) (U01DA050239 to F.T.; F31DA056226 to J.L.Z.; U01DA043799 to O.G., P50DA037844 to A.A.P., R01GM140287 to P.M.). G.d.G. was supported by the Brain and Behavior Research Foundation 2020 Young Investigator Award and M.K. by the TDRP (T31KT1859 UC) grant. This publication includes data generated at the UC San Diego IGM Genomics Center utilizing an Illumina NovaSeq 6000 that was purchased with funding from the NIH (S10OD026929).

## Author information

### Contributions

F.T. designed and coordinated the study. G.M. designed the overall bioinformatics analysis flow. J.L.Z. conducted bioinformatic analysis and data interpretation with inputs from F.T., G.M., and A.A.P.. A.J.H. conducted ChromVar analysis. O.G. and G.d.G. designed the behavioral protocol for cocaine IVSA. G.d.G. performed the behavioral experiments. M.K. performed the electrophysiological experiments. L.L.C. dissected the brain samples. H.L. performed snATAC-seq experiments. N.P. performed the snRNAseq experiments for the CeA and BLA. A.A.P., D.M. and P.M. coordinated and executed the gene expression predictions for naive HS rats using eQTL data. A.S.C. provided genotype data for exploratory analysis. J.L.Z., A.P., G.M. and F.T. wrote the manuscript with contributions from all authors.

### Corresponding authors

Correspondence to Graham McVicker or Francesca Telese.

