## Supplemental Information for "Cocaine addiction-like behaviors are associated with long-term changes in gene regulation, energy metabolism, and GABAergic inhibition within the amygdala"

### Supplementary Information

#### Supplementary Figures

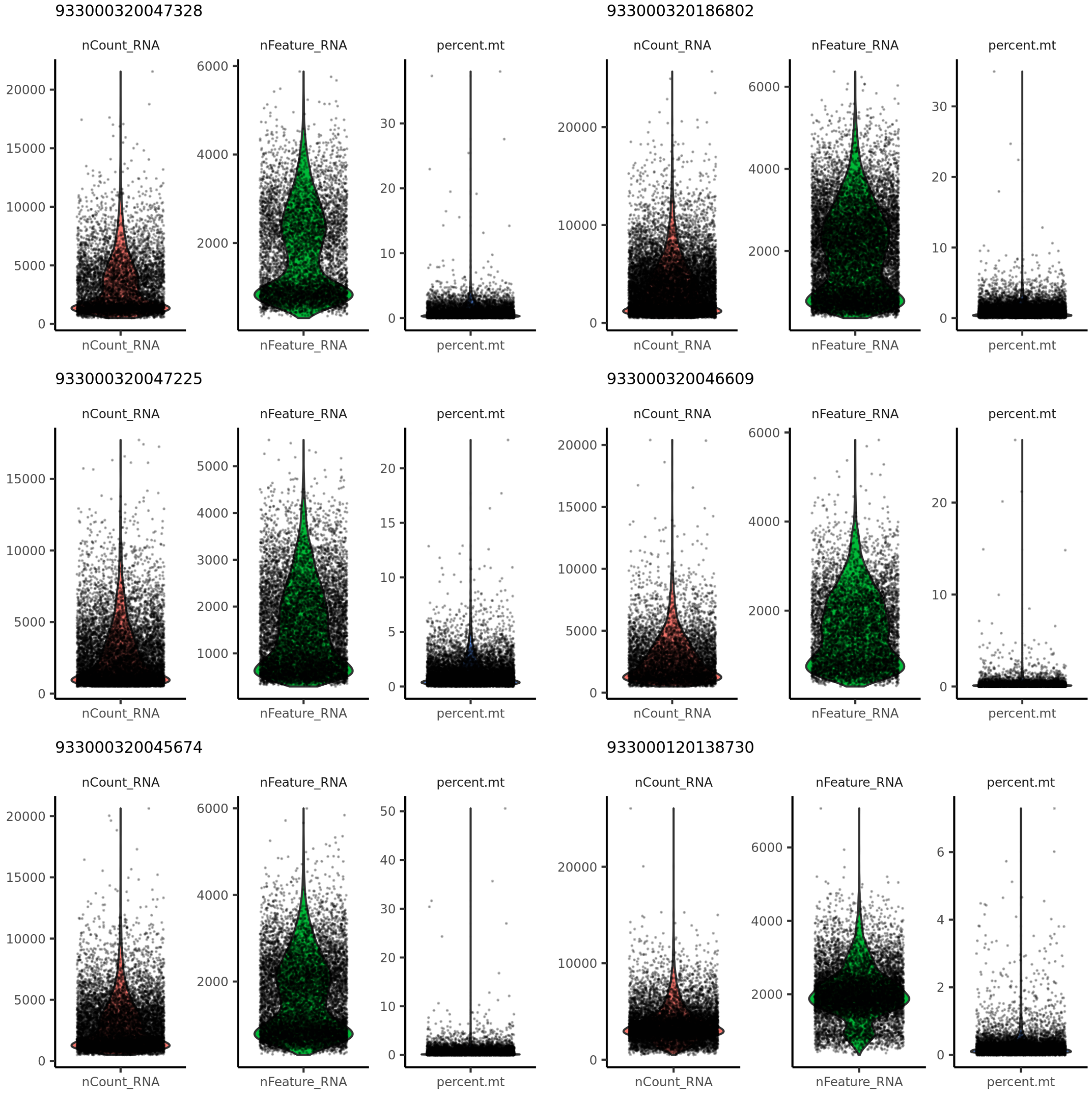


**Supplementary Figure 1.** snRNA-seq profiles of 6 high AI rats. For each rat the number of unique genes detected per cell (nFeature_RNA), total number of reads within each cell (nCount_RNA), and percentage of percent mitochondrial reads are shown for each cell. Quality metrics calculated with Seurat.

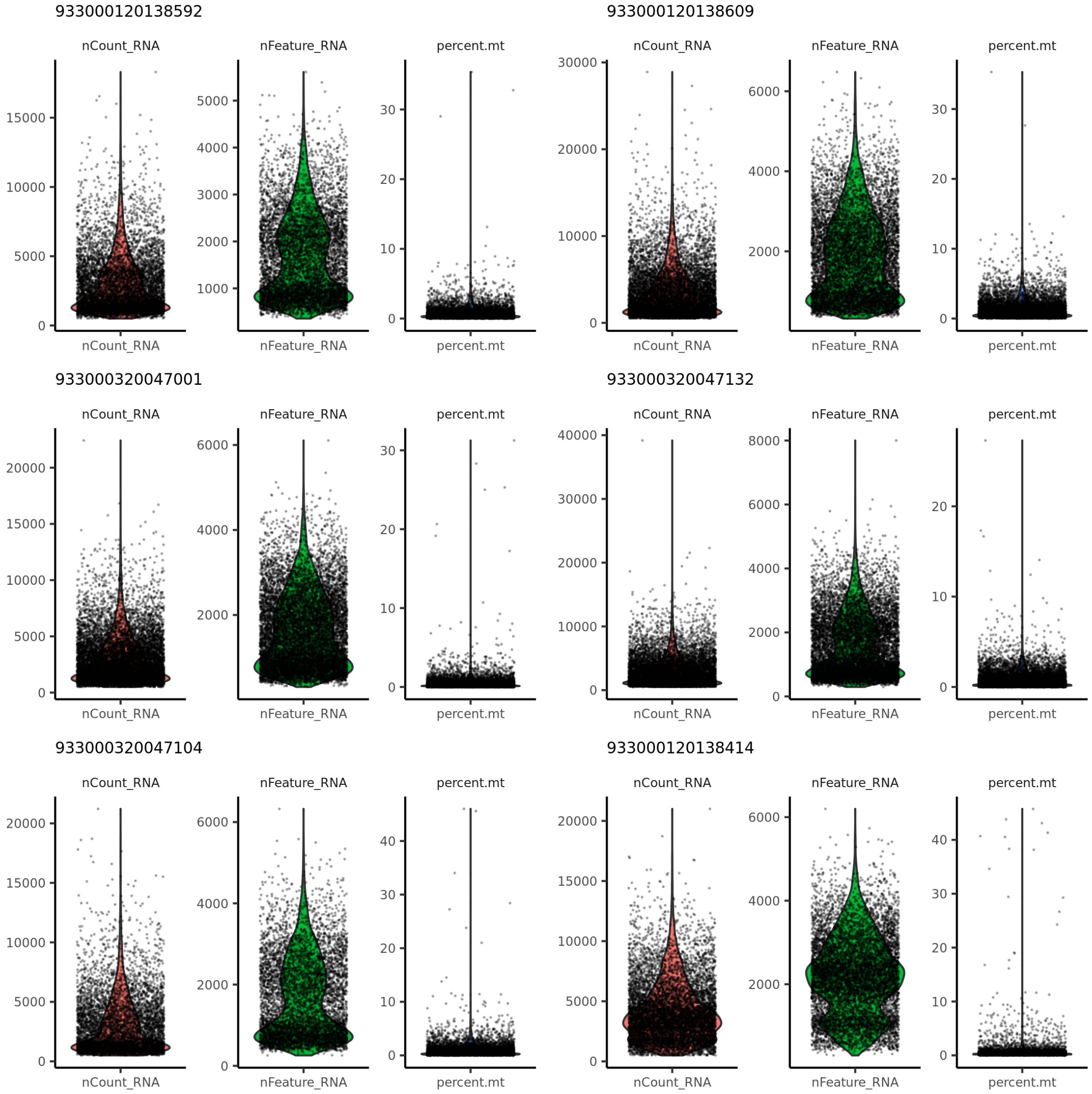


**Supplementary Figure 2.** snRNA-seq profiles of 6 low AI rats. For each rat the number of unique genes detected per cell (nFeature_RNA), total number of reads within each cell (nCount_RNA), and percentage of percent mitochondrial reads are shown for each cell. Quality metrics calculated with Seurat.

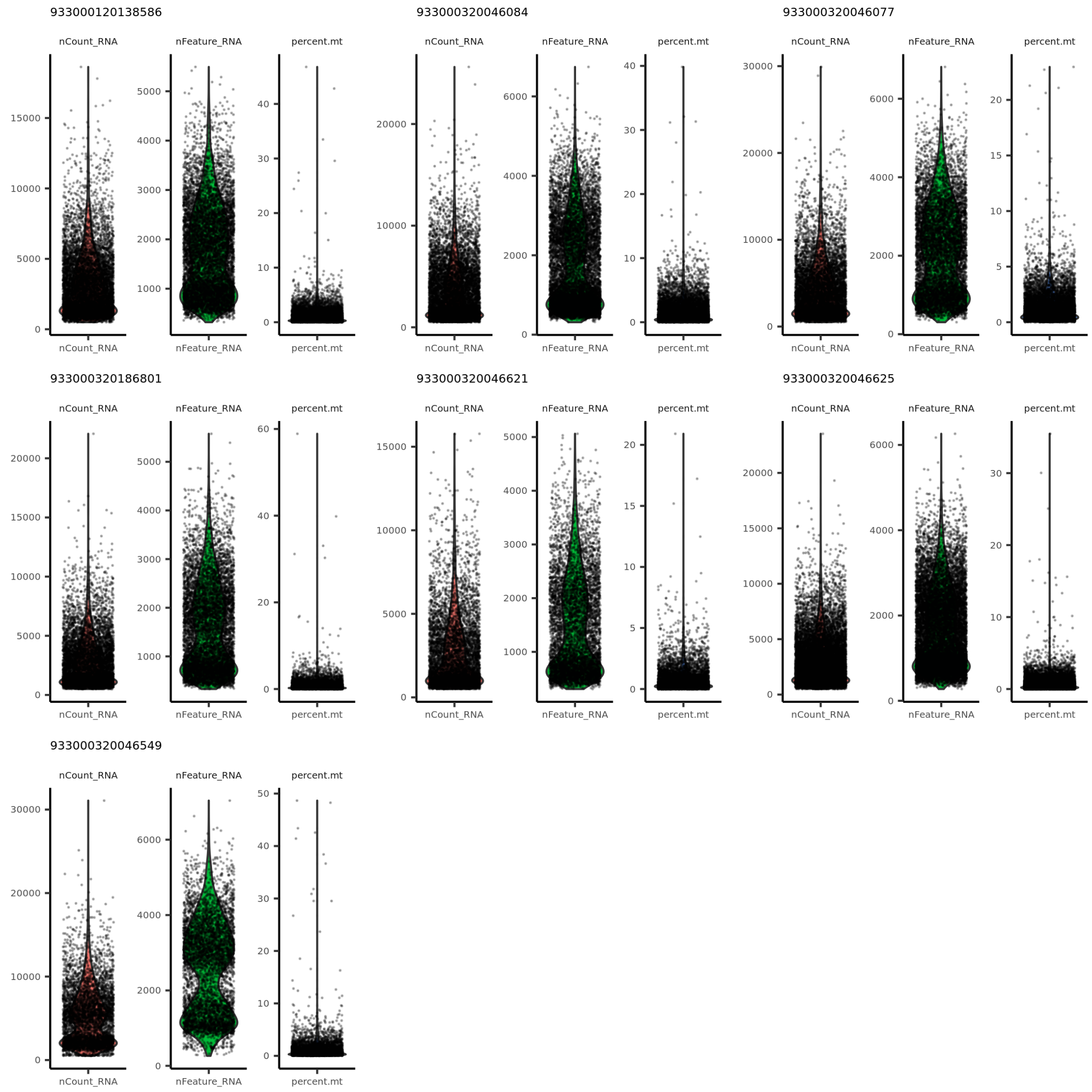


**Supplementary Figure 3.** snRNA-seq profiles of 7 naive rats. For each rat the number of unique genes detected per cell (nFeature_RNA), total number of reads within each cell (nCount_RNA), and percentage of percent mitochondrial reads are shown for each cell. Quality metrics calculated with Seurat.

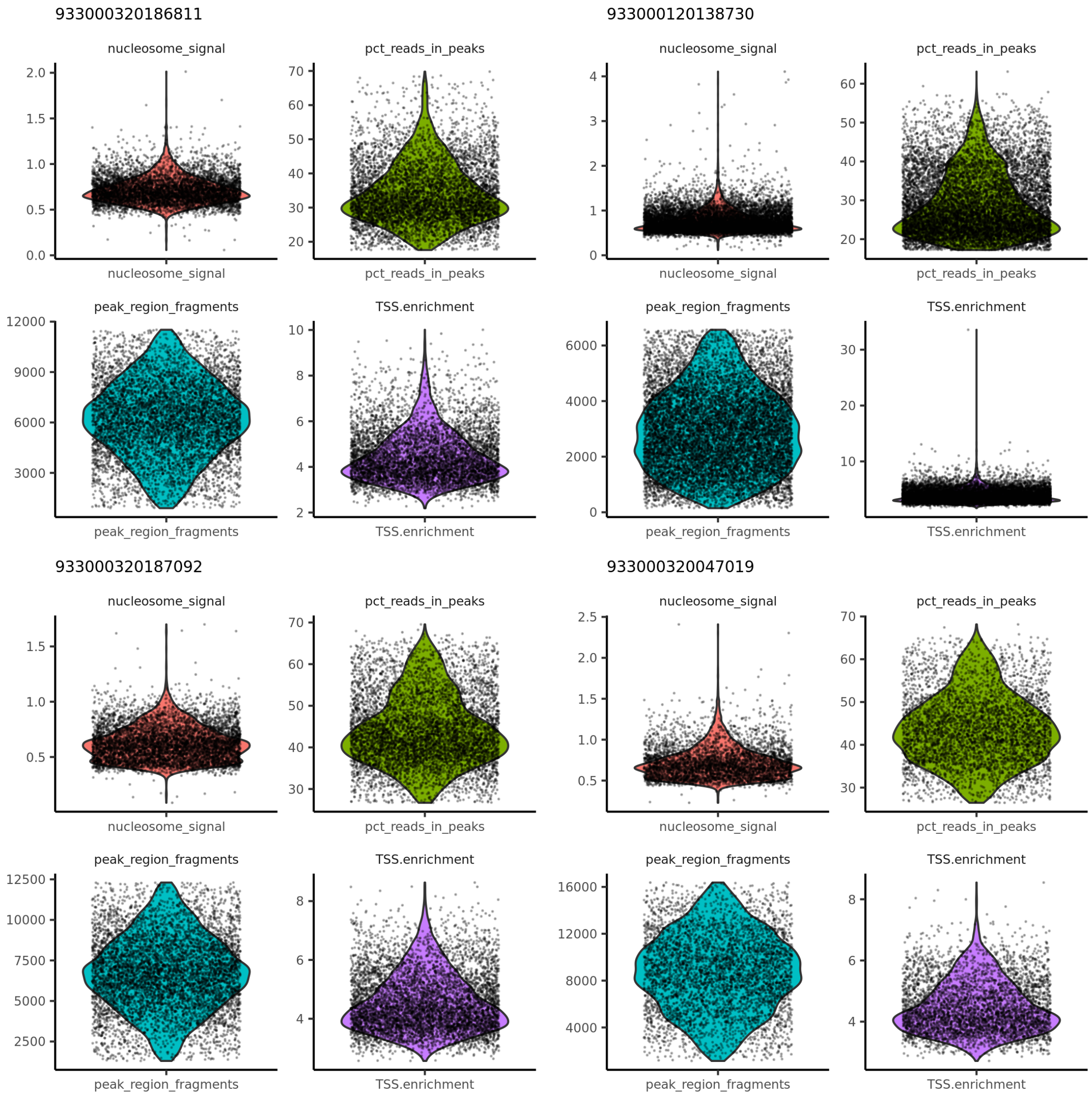


**Supplementary Figure 4.** snATAC-seq profiles of 4 high AI rats. For each rat the ratio of mononucleosomal to nucleosome-free fragments (nucleosome_signal), percentage of fragments that fall within ATAC-seq peaks (pct_reads_in_peaks), total number of fragments in peaks (peak_region_fragments), and transcription start site enrichment score (TSS.enrichment) are shown for each cell. Quality metrics calculated with Signac.

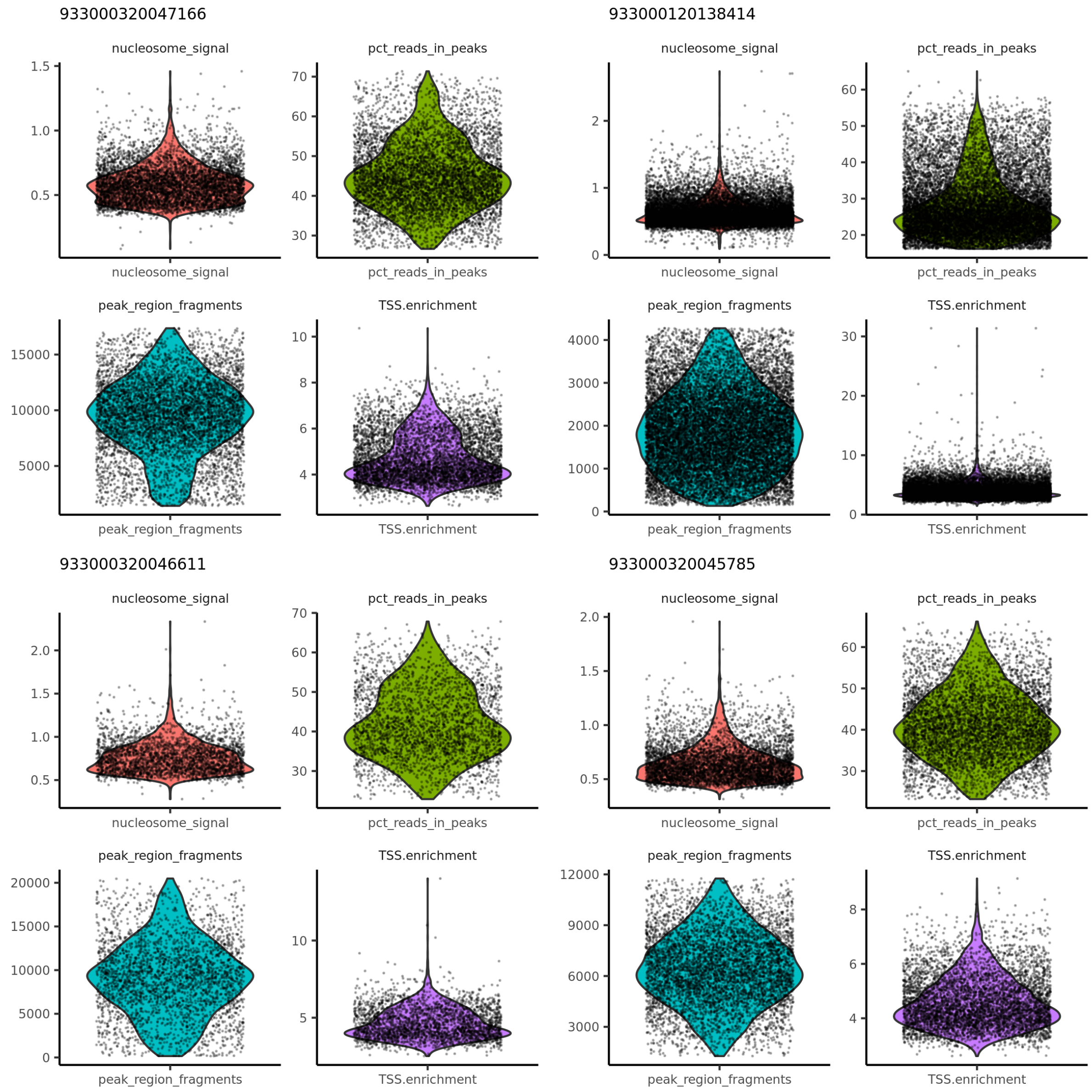


**Supplementary Figure 5.** snATAC-seq profiles of 4 low AI rats. For each rat the ratio of mononucleosomal to nucleosome-free fragments (nucleosome_signal), percentage of fragments that fall within ATAC-seq peaks (pct_reads_in_peaks), total number of fragments in peaks (peak_region_fragments), and transcription start site enrichment score (TSS.enrichment) are shown for each cell. Quality metrics calculated with Signac.

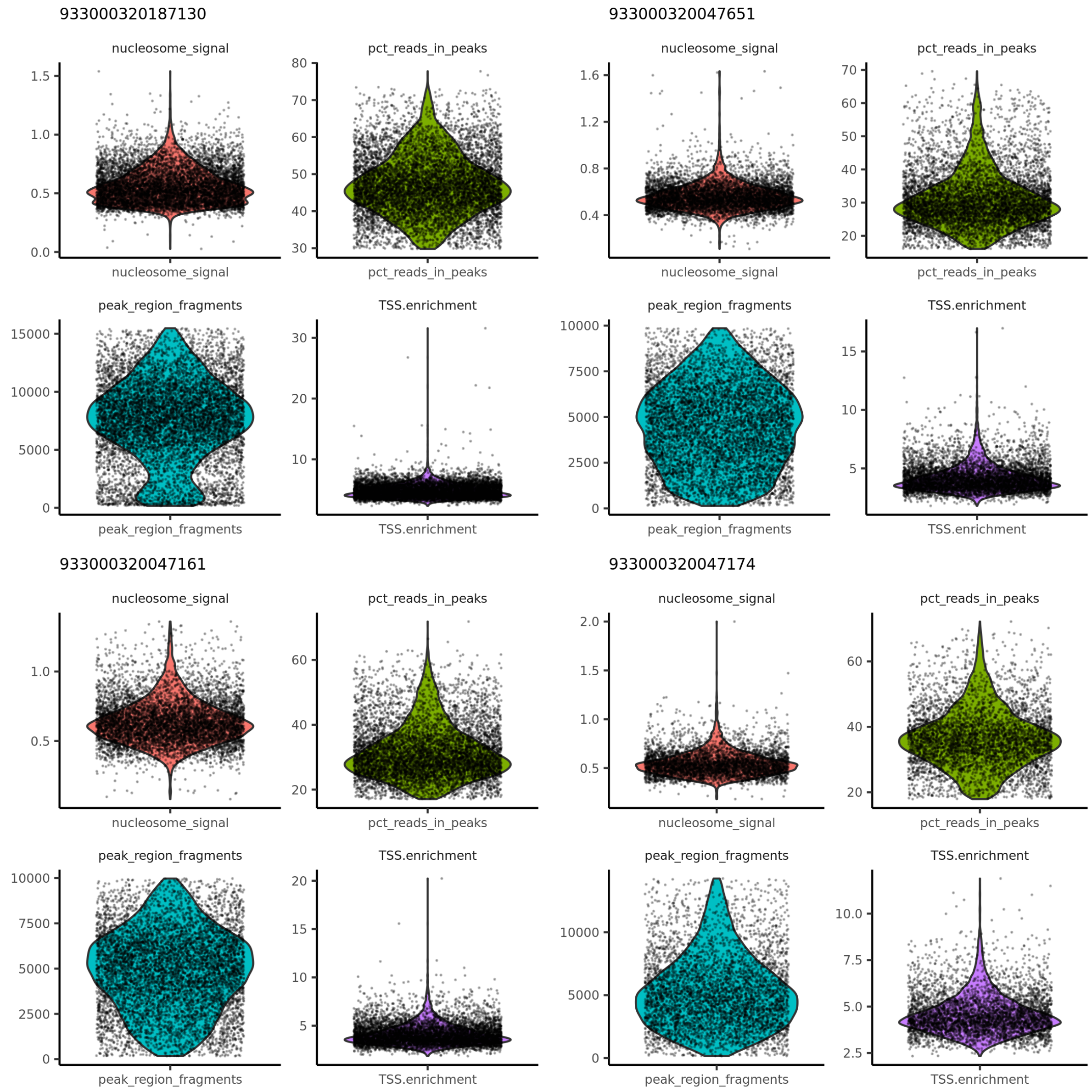


**Supplementary Figure 6.** snATAC-seq profiles of 4 low naive rats. For each rat the ratio of mononucleosomal to nucleosome-free fragments (nucleosome_signal), percentage of fragments that fall within ATAC-seq peaks (pct_reads_in_peaks), total number of fragments in peaks (peak_region_fragments), and transcription start site enrichment score (TSS.enrichment) are shown for each cell. Quality metrics calculated with Signac.

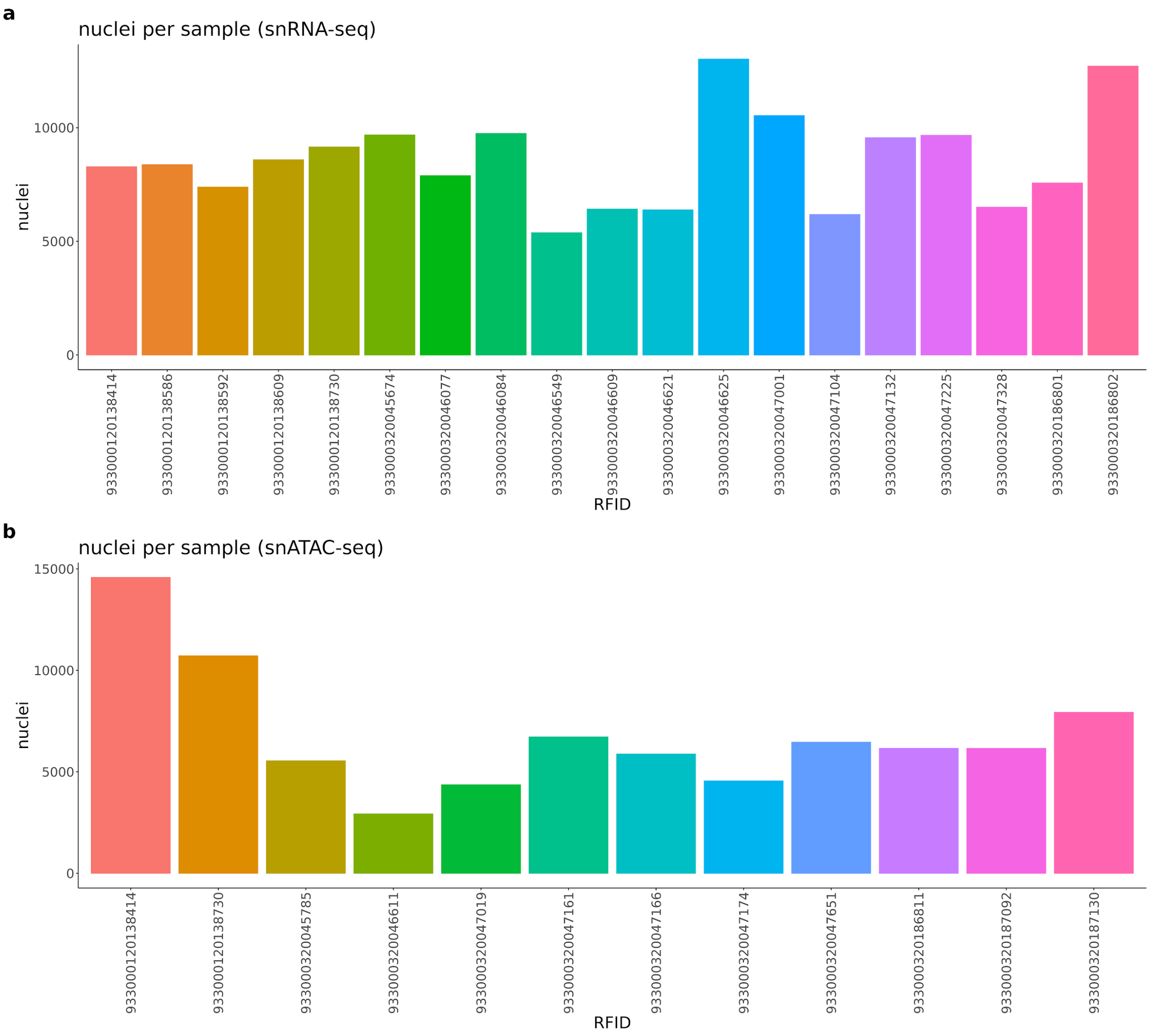


**Supplementary Figure 7.** Number of nuclei coming from each sample in the a) snRNA-seq and b) snATAC-seq datasets. Mean nuclei per sample is a) 8579 in the snRNA-seq, and b) 6826 in the snRNA-seq.

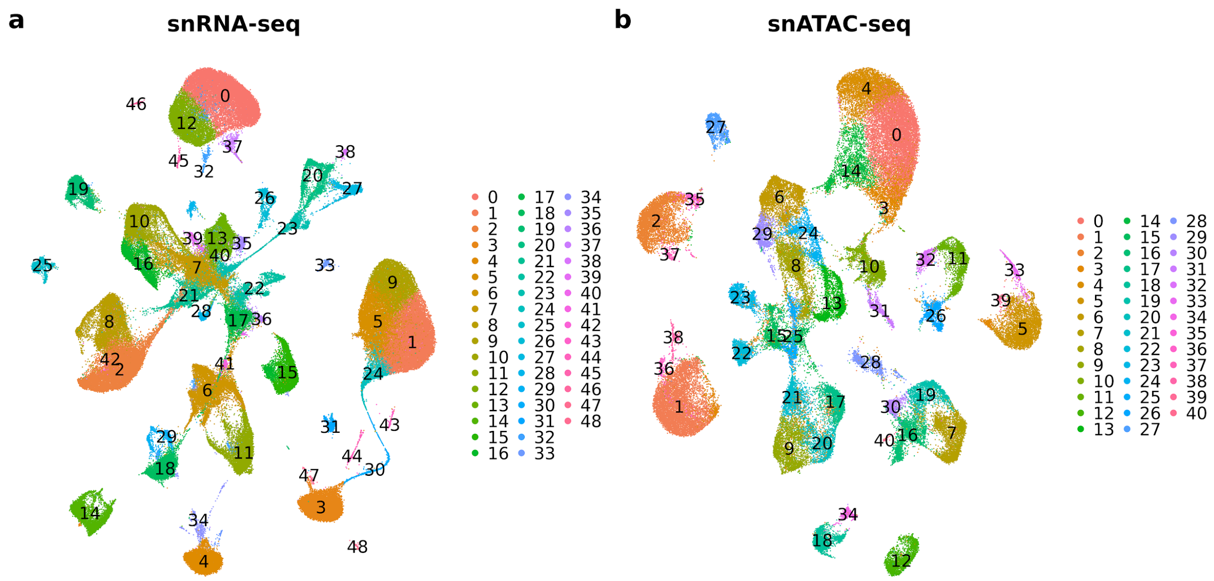


**Supplementary Figure 8.** UMAP visualization of the clusters identified in integrated single-cell data sets. (**a**) Clustering of integrated snRNA-seq dataset revealed 49 clusters. We first performed a k-nearest neighbors analysis (KNN) using the first 30 dimensions calculated by reciprocal principal component analysis (PCA). This was implemented with the FindNeighbors() function in Seurat. Next we used a modularity optimization technique using the Louvain algorithm to cluster the data, implemented with the FindClusters() function in Seurat with a resolution parameter of 0.8. (**b**) Clustering of integrated snATAC-seq data revealed 41 clusters. Latent semantic indexing (LSI) was used for dimensionality reduction rather than PCA. The first 30 dimensions minus the first dimension were used for KNN and clustering and the algorithm used for clustering was the smart local moving (SLM) algorithm. These steps were implemented with the same Seurat functions.

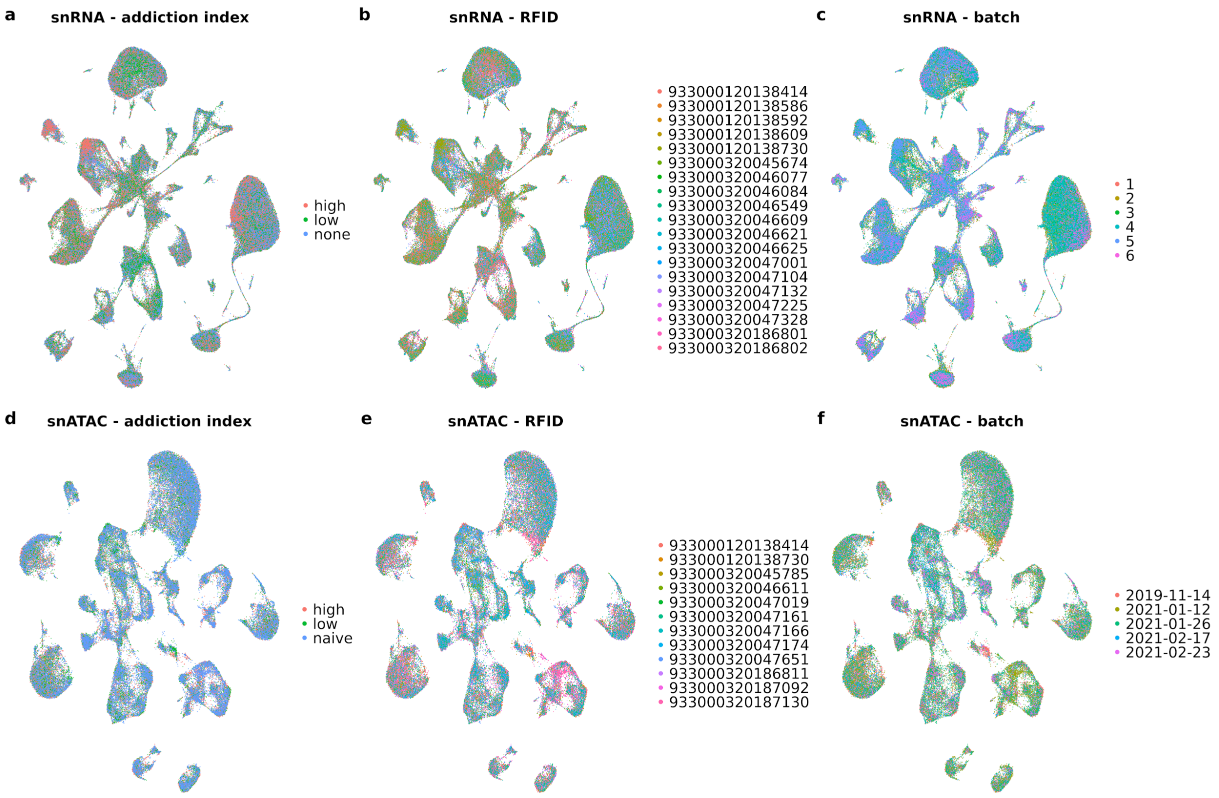


**Supplementary Figure 9.** UMAPS of snRNA-seq and snATAC-seq profiles, respectively, following batch correction of integrated datasets, grouped on: addiction index (a, d), rat sample (b, e), and batch information (c, f). These plots demonstrate that cells do not cluster by any of these covariates following batch correction. Integration and batch correction of the snRNA-seq dataset was performed using SCTransform while Harmony was used for the snATAC-seq dataset.

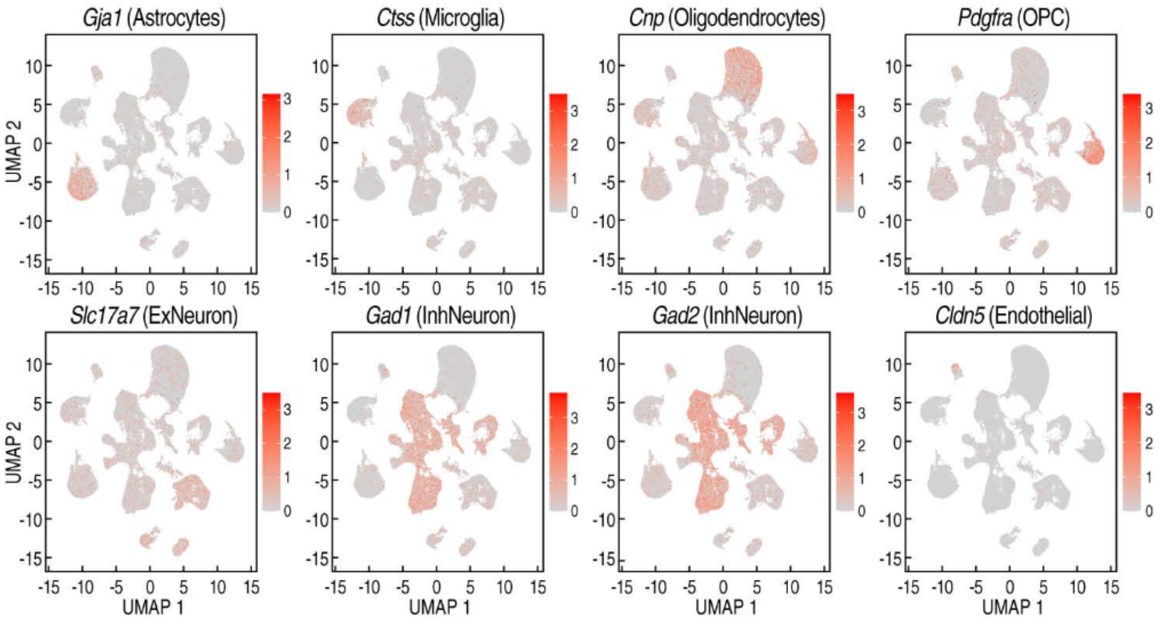


**Supplementary Figure 10.** Feature plots showing gene activity of marker genes for each major cell type in the snATAC-seq data. Gene activity was calculated with the \`GeneActivity()\` function in Signac. This quantifies the number of fragments mapping anywhere within a 2kb window of an annotated gene in the genome. The gene activity information was used for integration of the snATAC-seq dataset with the snRNA-seq dataset and for imputing gene expression into the cells of the snATAC-seq dataset (see Fig. 2d).

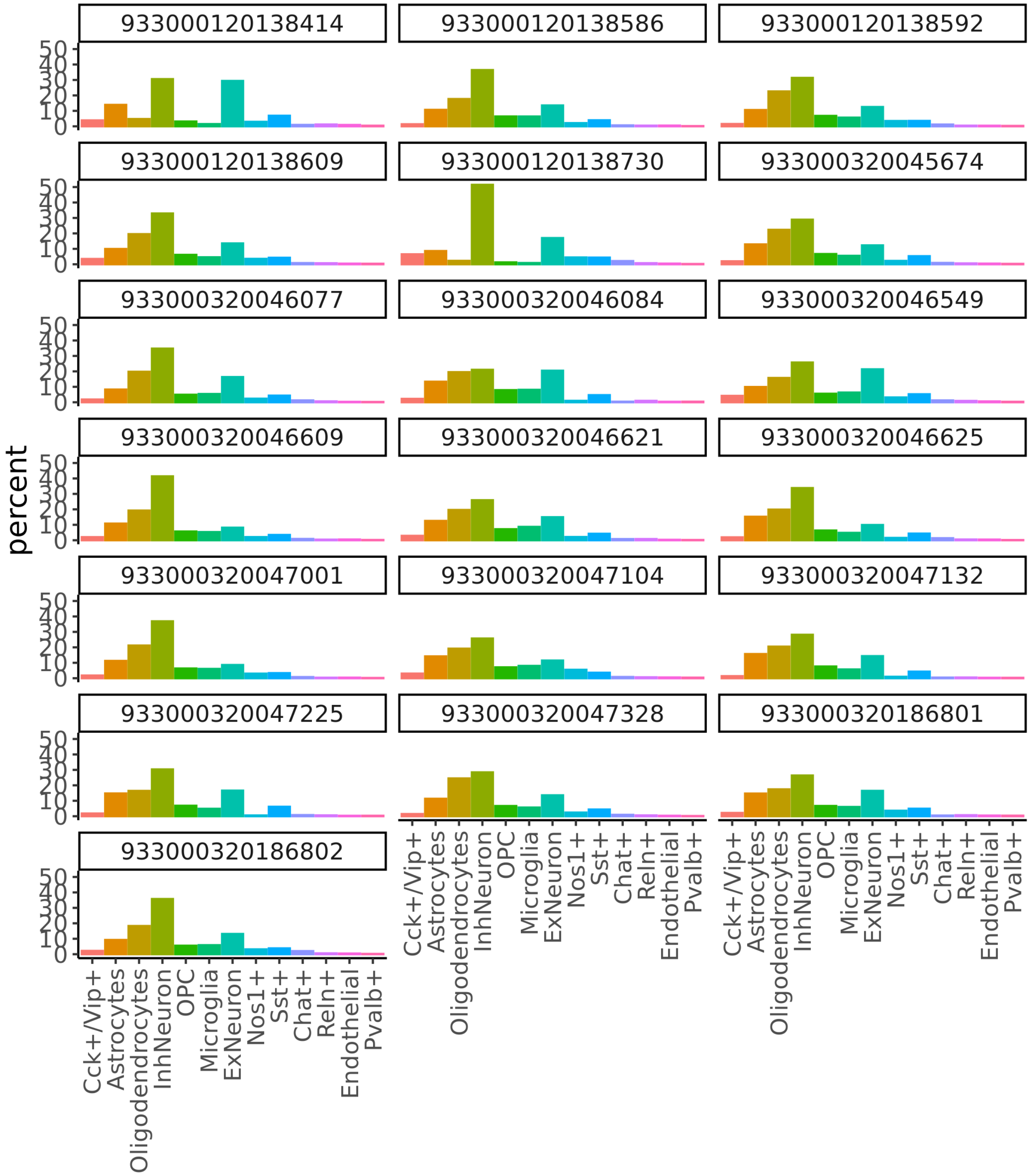


**Supplementary Figure 11.** Barplots show counts of each cell type within each sample, labeled by RFID, for the snRNA-seq samples.

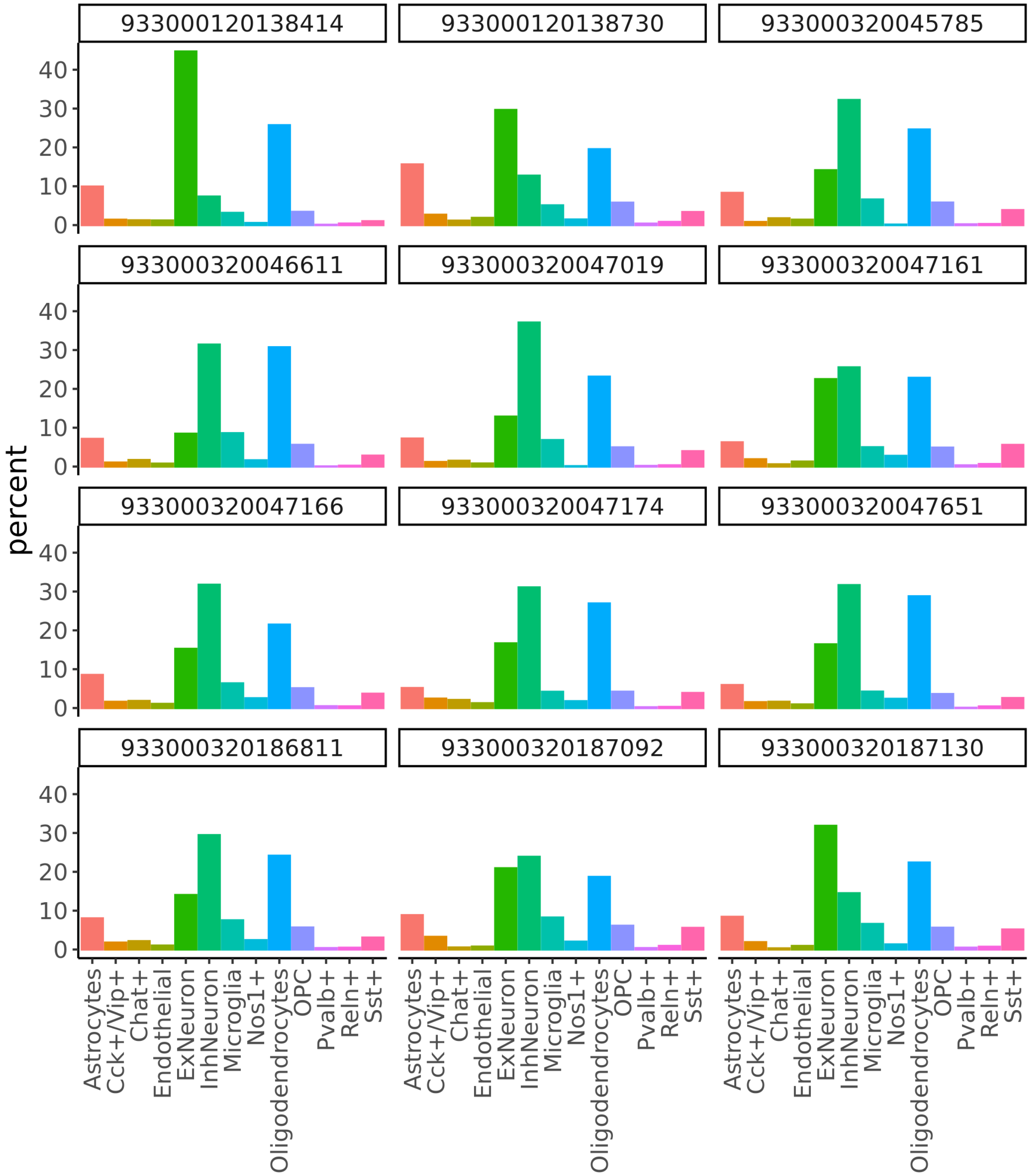


**Supplementary Figure 12.** Barplots show counts of each cell type within each sample, labeled by RFID, for the snATAC-seq samples.

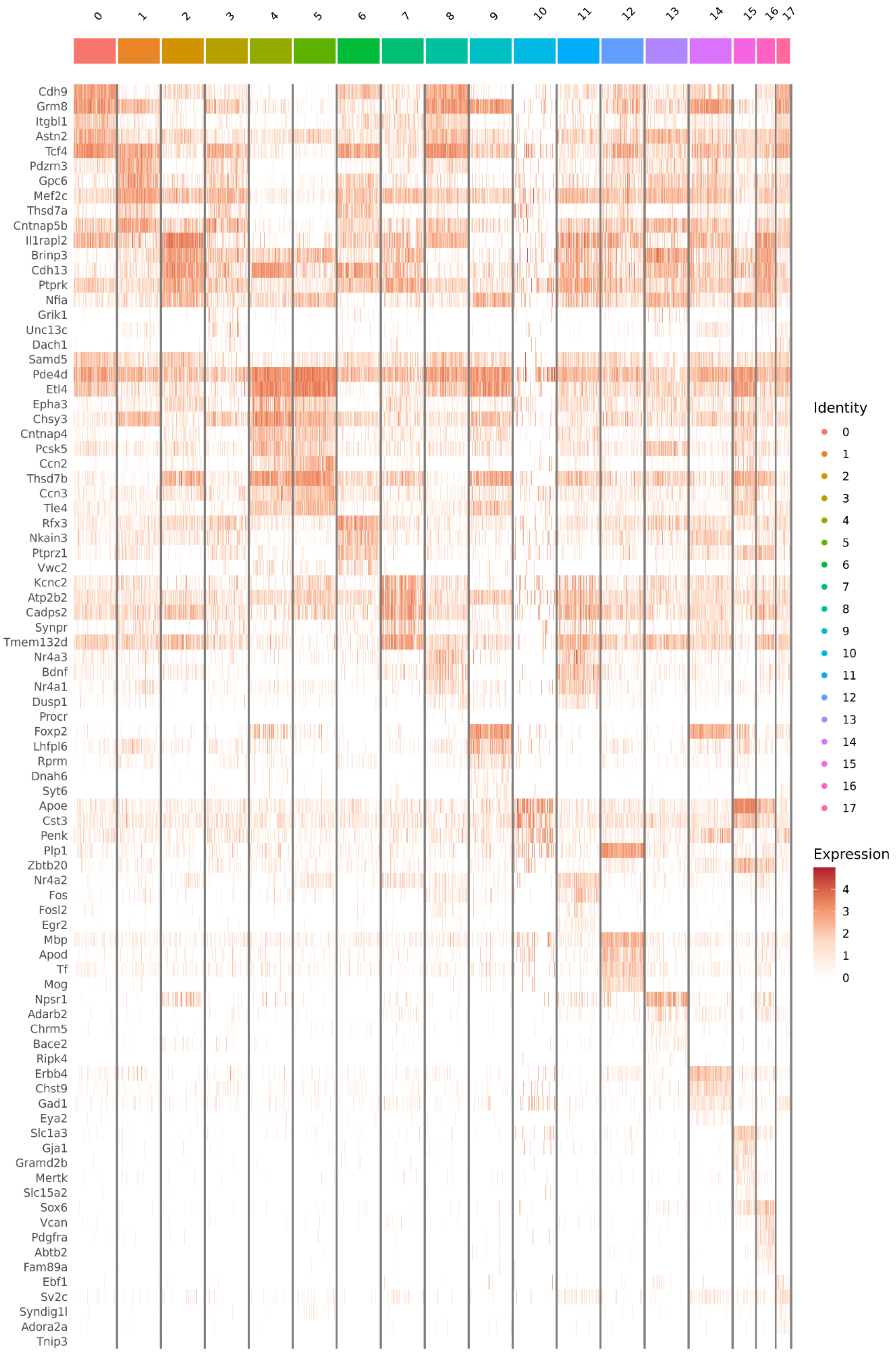


**Supplementary Figure 13.** Heatmap of top five marker gene expression within subclustered excitatory
neurons.

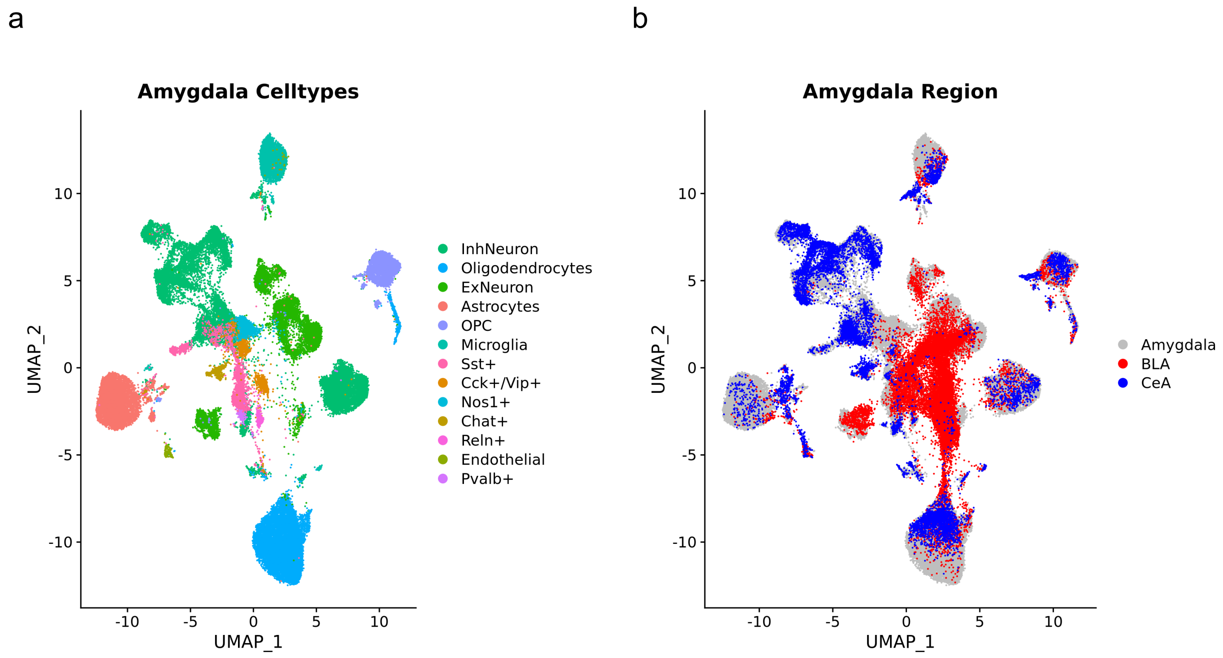


**Supplementary Figure 14.** Co-clustering of snRNA-seq from a CEA sample, a BLA sample, and the whole amygdala samples from all of the naive rats in our study. a) UMAP with cells colored by cell type cluster assignments. b) UMAP with cells colored by source tissue, where "Amygdala" refers to the set of all amygdala samples from the naive rats in our study.

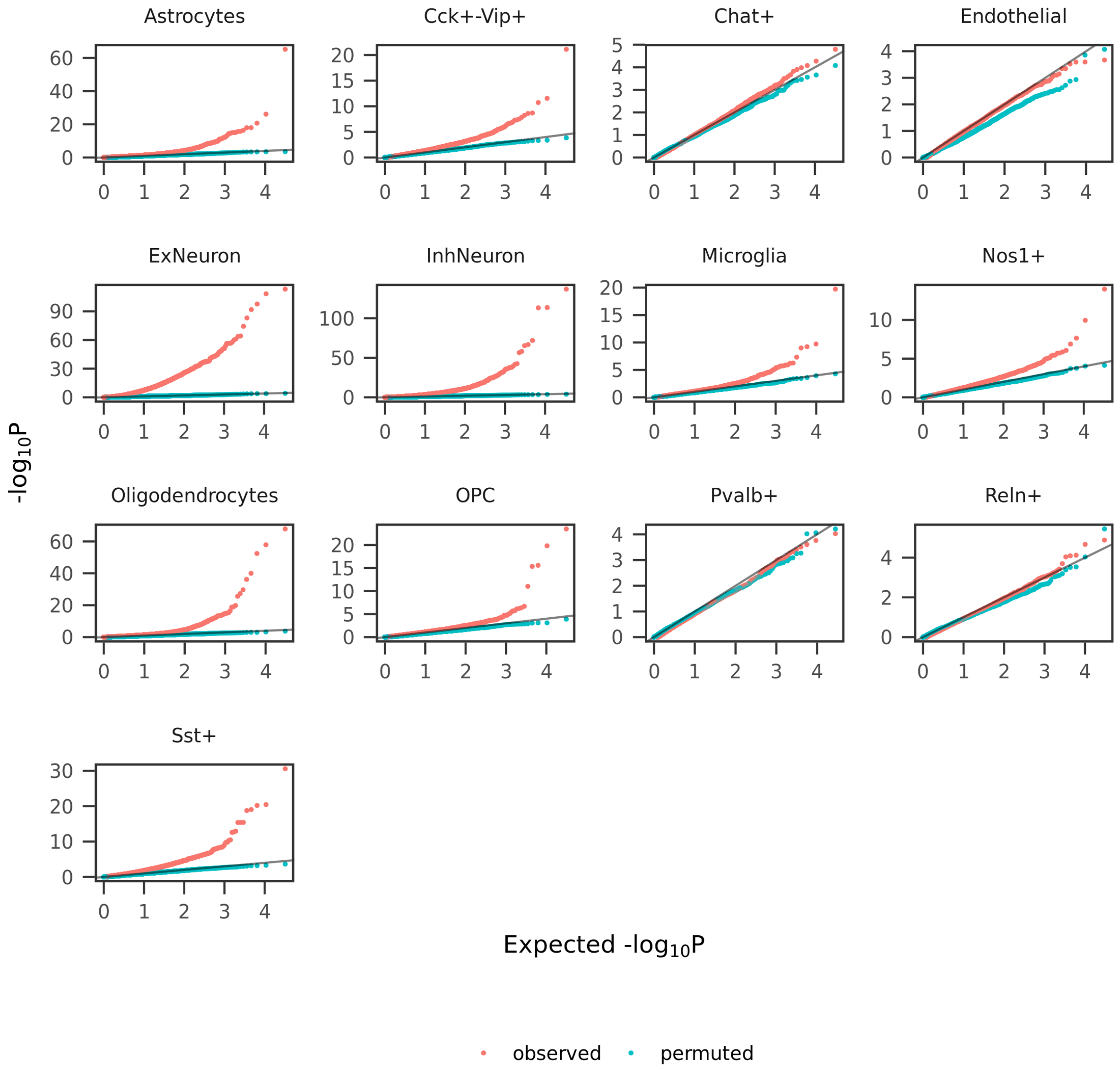


**Supplementary Figure 14.** QQ plots showing distribution of p-values for our differential gene expression analysis performed on our observed versus permuted data (AI labels associated with each cell were shuffled). The negative binomial test was the statistical test used for the analysis of both the observed and permuted datasets.

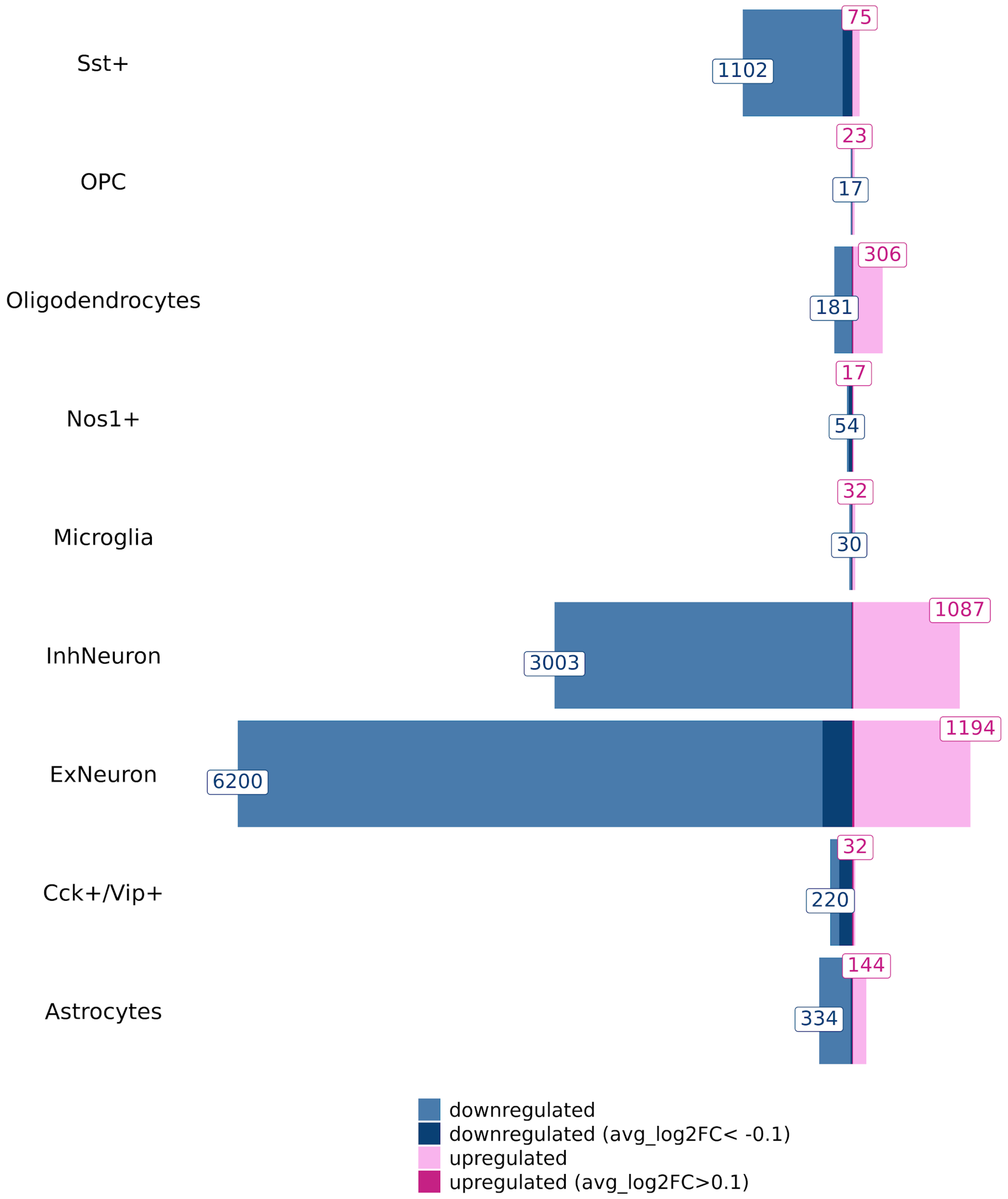


**Supplementary Figure 16.** Barplot showing numbers (labeled) of significant (FDR<10%) up- and downregulated DEGs by cell type. Darker shades indicate DEGs with a large fold change (abs(avg_$\log_2$FC)≥0.1).

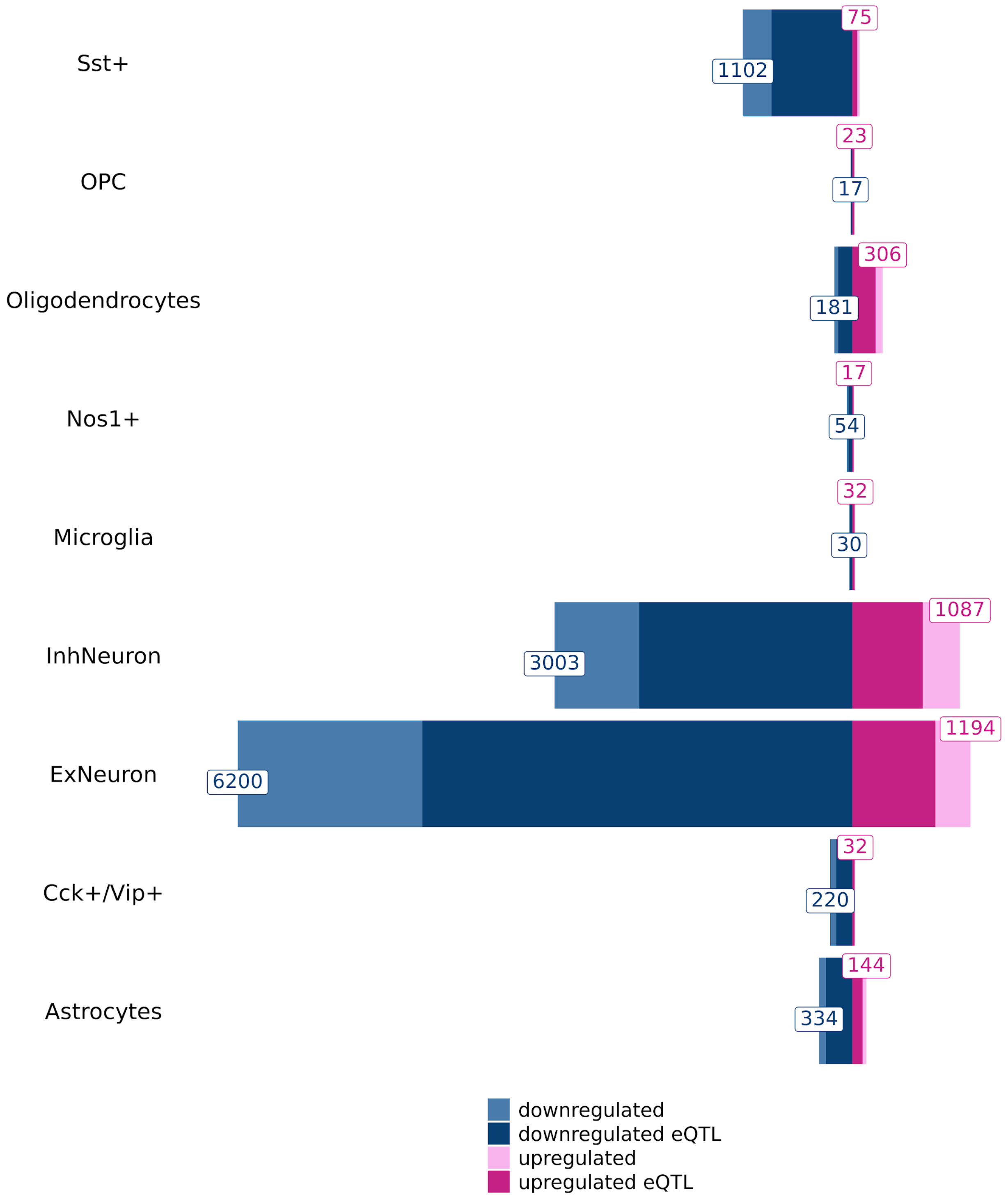


**Supplementary Figure 17.** Barplot showing numbers (labeled) of significant (FDR<10%) up- and downregulated DEGs by cell type. Darker shades indicate DEGs that are also significant eQTLs in rat brain tissues.

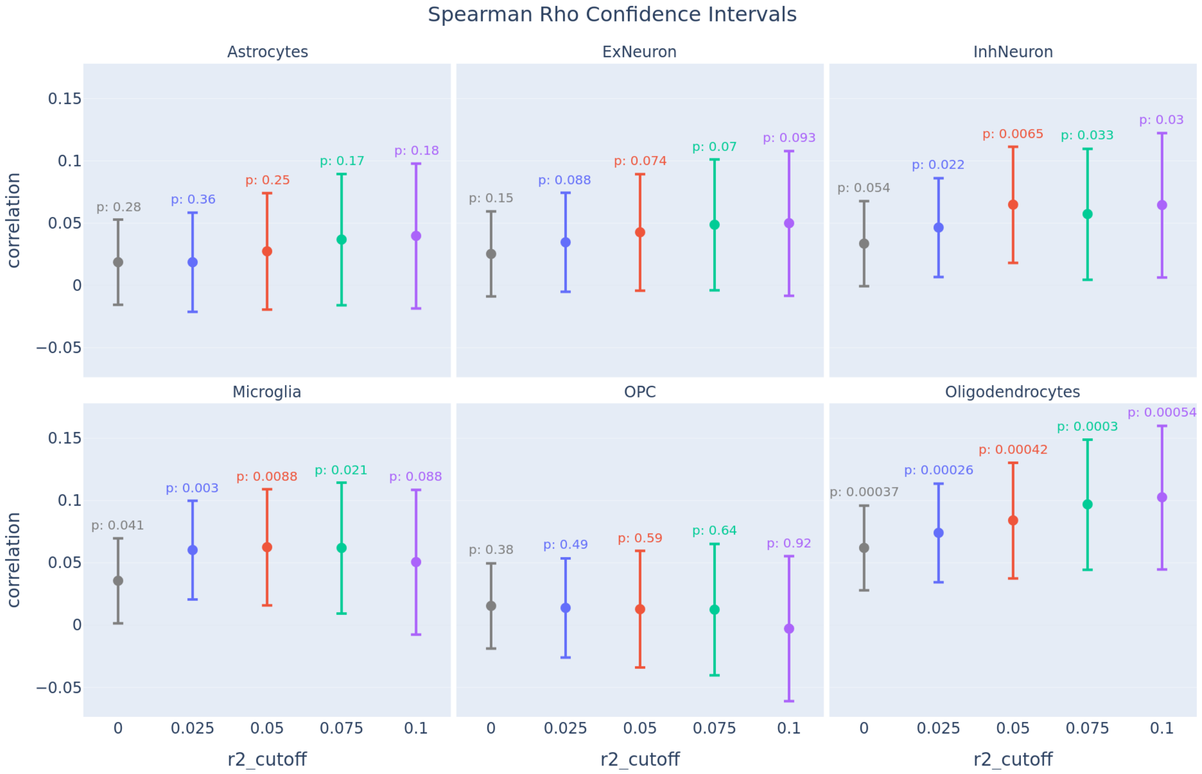


**Supplementary Figure 18.** Spearman's Rho (ρ) confidence intervals for the correlations between predicted vs. observed differences in gene expression (high vs. low AI) for each major cell type. We filtered the genes in the predicted set based on the $r^2$ metric, or predictive accuracy, of their predictive models. Each color corresponds to a different cutoff for $r^2$. We see significant correlations (p<0.05) in microglia, oligodendrocytes and inhibitory neurons. We also observe a general trend of increasing correlation coefficients ρ as we increase the $r^2$ cutoff. Spearman's correlation coefficient ρ is plotted on the y-axis; $r^2$ cutoff is plotted on the x-axis.

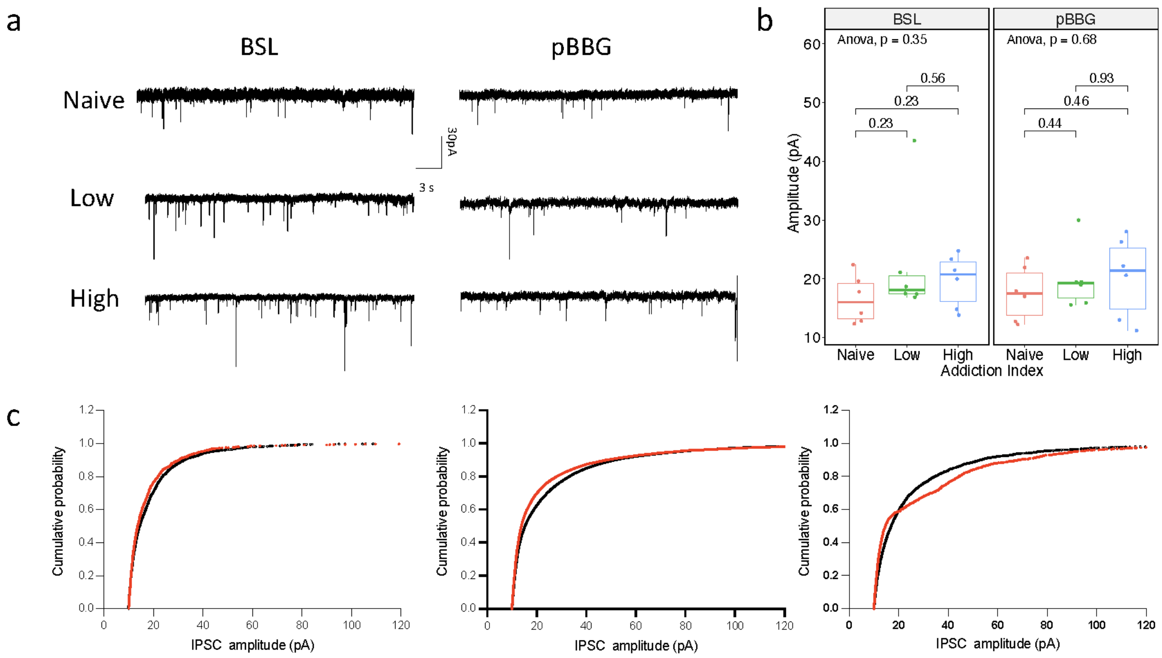


**Supplementary Figure 19.** Summary of electrophysiology experiments studying GABA transmission in the central amygdala. a) Representative traces of sIPSC frequencies for baseline (BSL) and following treatment with pBBG (pBBG) in naive, low AI and high AI rats. b) ANOVA test comparing mean amplitude in BSL vs. pBBG across naive, low AI and high AI rats (degrees of freedom = 2). c) Cumulative probability plots of the peak amplitude for naive, low and high rats.

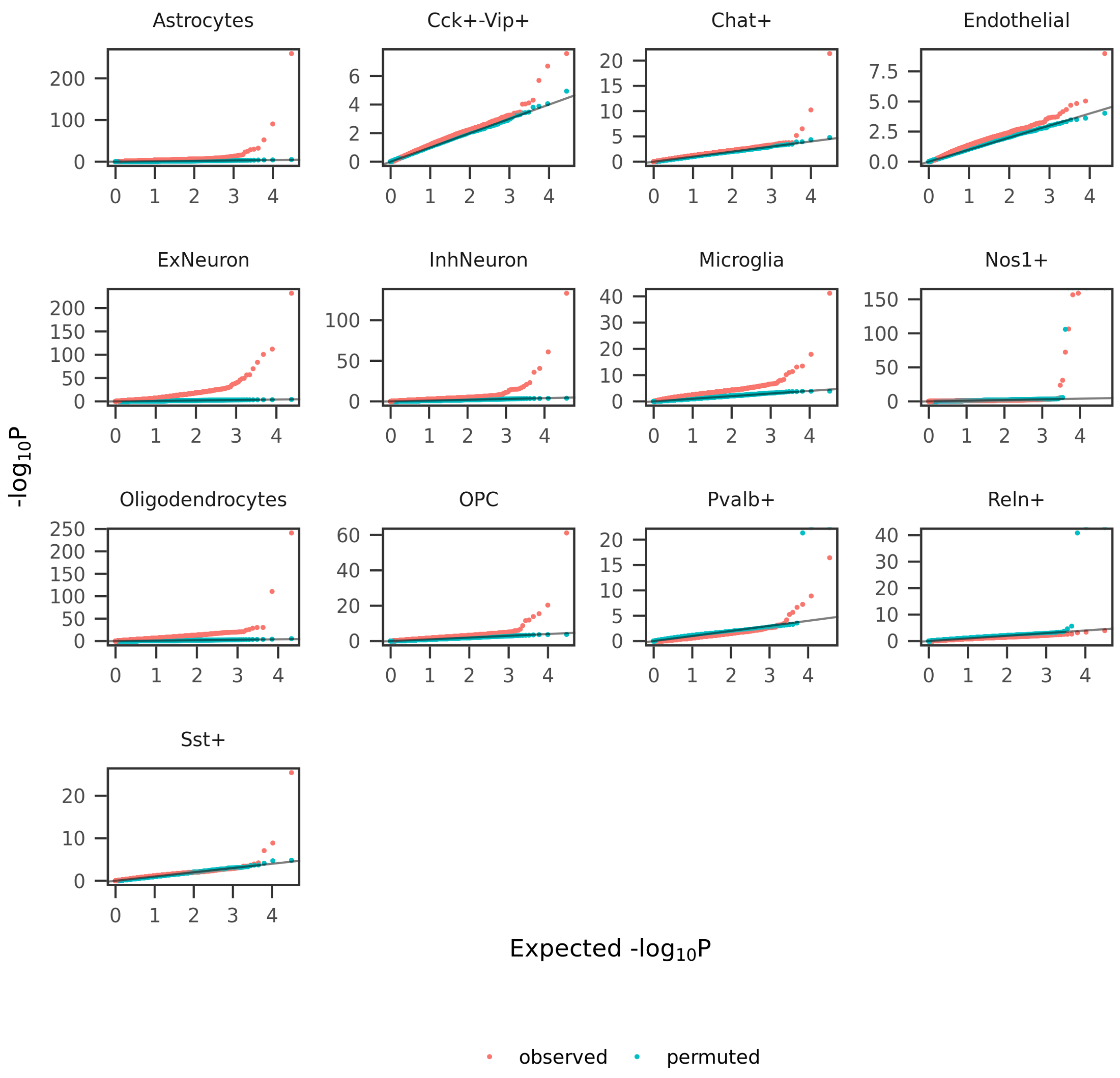


**Supplementary Figure 20.** QQ plots showing distribution of p-values for our differential peak accessibility analysis performed on our observed versus permuted data (AI labels associated with each cell were shuffled). The negative binomial test was used for the analysis of both the observed and permuted datasets.

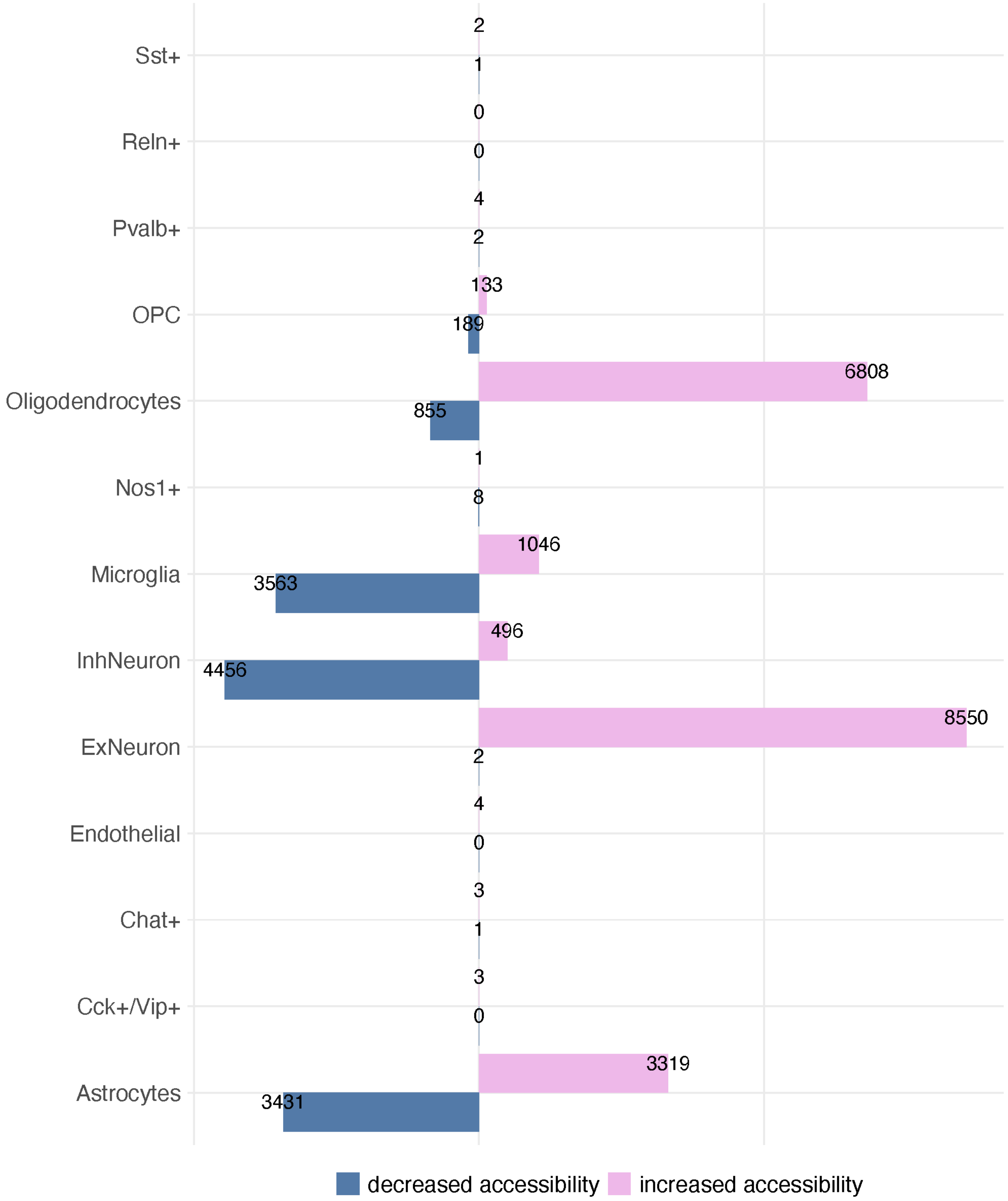


**Supplementary Figure 21.** Bar plot showing number of significant (FDR<10%) differentially accessible peaks between high vs. low rats in each cell type.

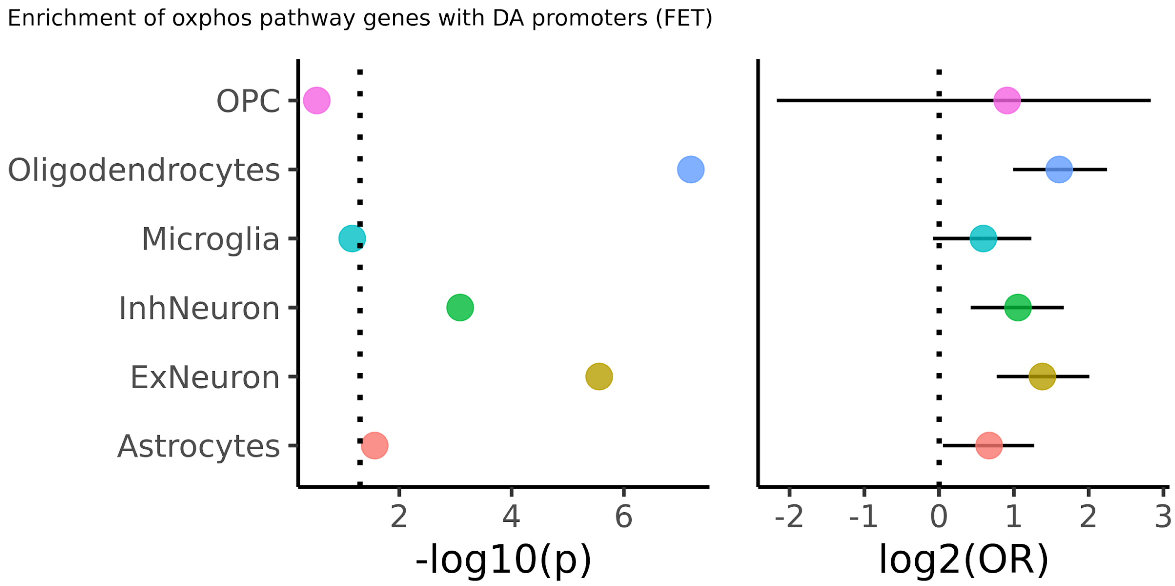


**Supplementary Figure 22.** Dot plots showing results of Fisher's Exact Test measuring enrichment of differentially accessible promoter regions (FDR<10%) in DEGs (FDR<10%) compared to non-DEGs for genes belonging to the oxidative phosphorylation pathway (as defined by the KEGG database).

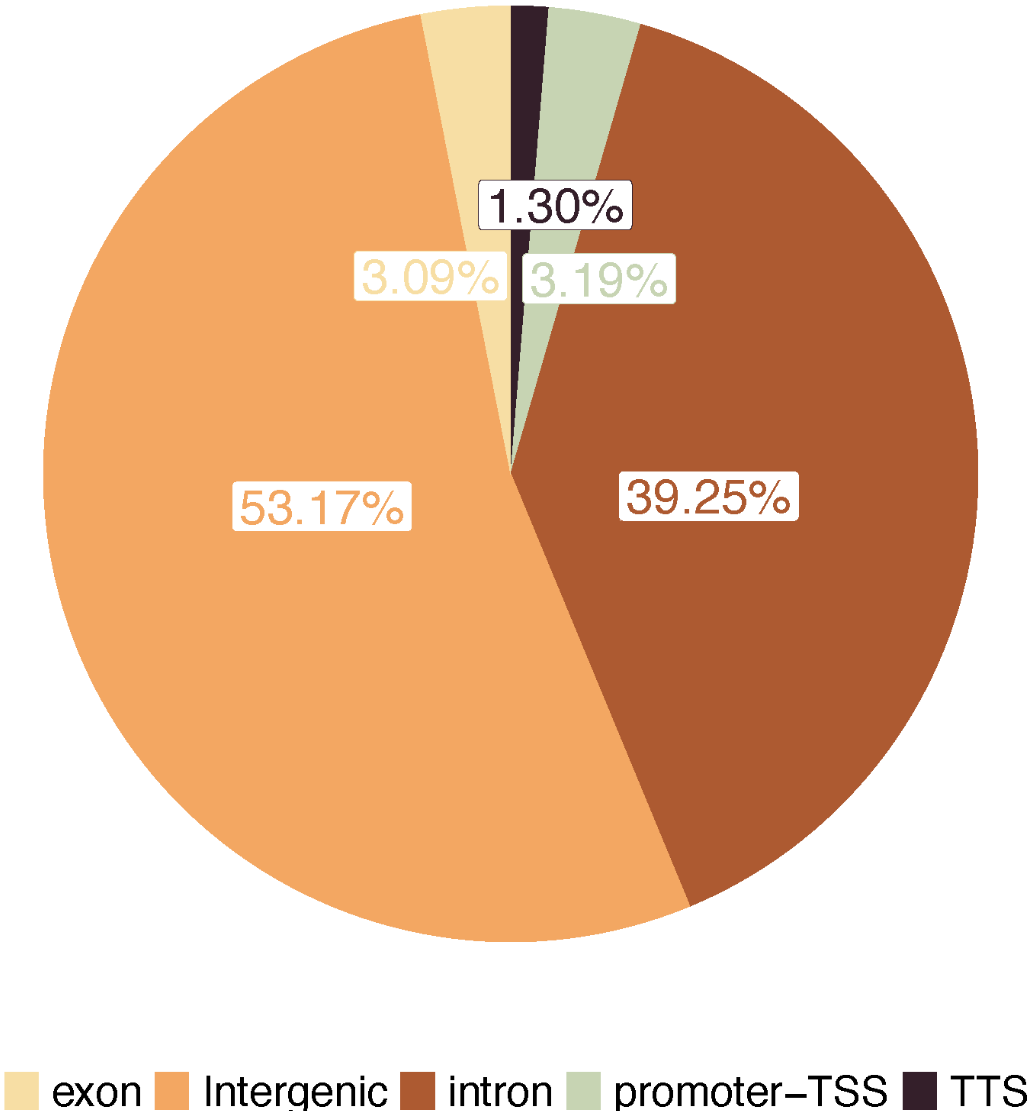


**Supplementary Figure 23.** Pie chart showing genomic annotations of all OCRs in our snATAC-seq dataset across all rats.

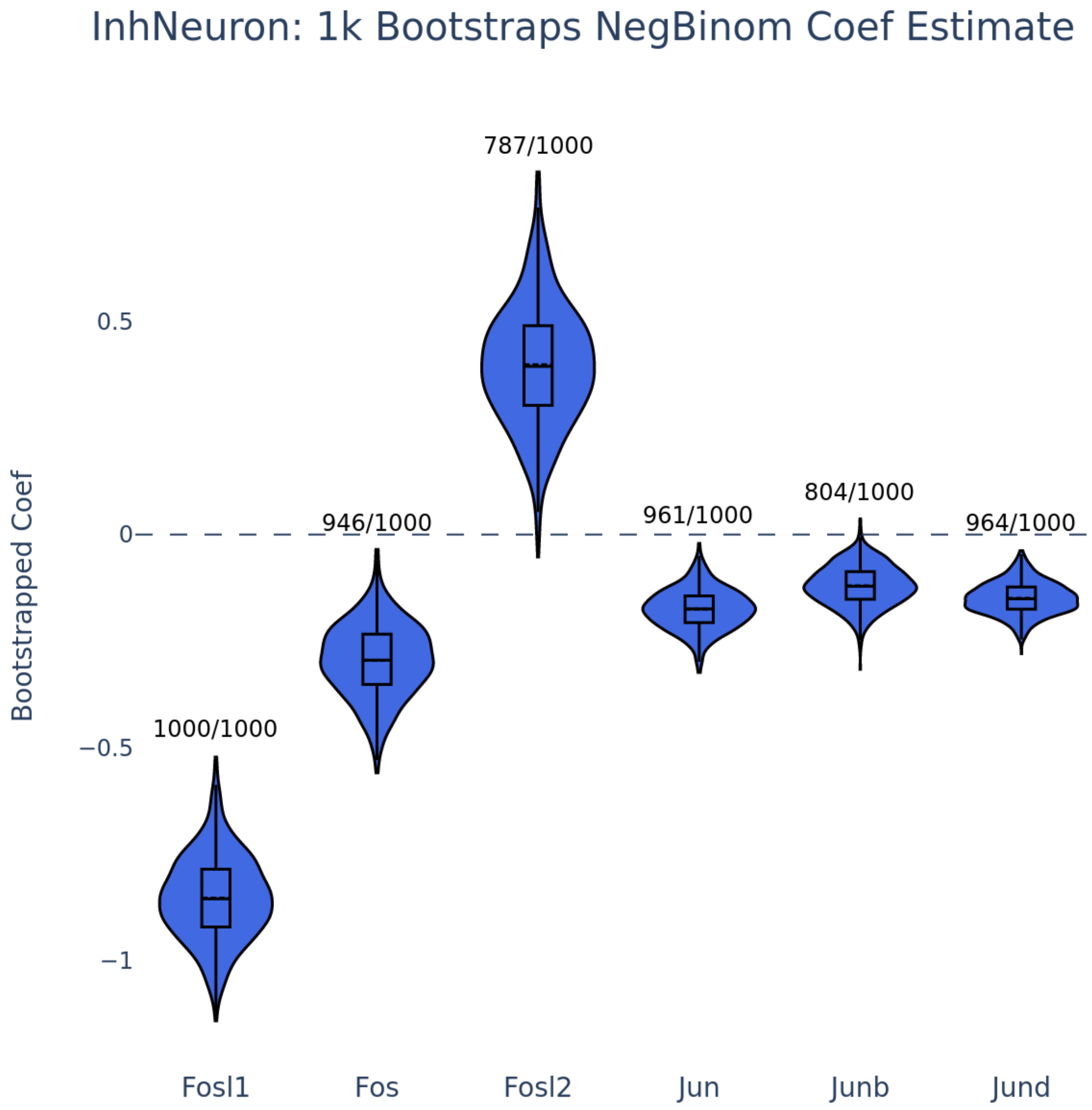


**Supplementary Figure 24.** Violin plots showing DEG analysis in InhNeurons over 1000 bootstrap iterations. Each violin shows the distribution of log2FC results per iteration. The fraction represents the number of significant iterations (FDR<10%).

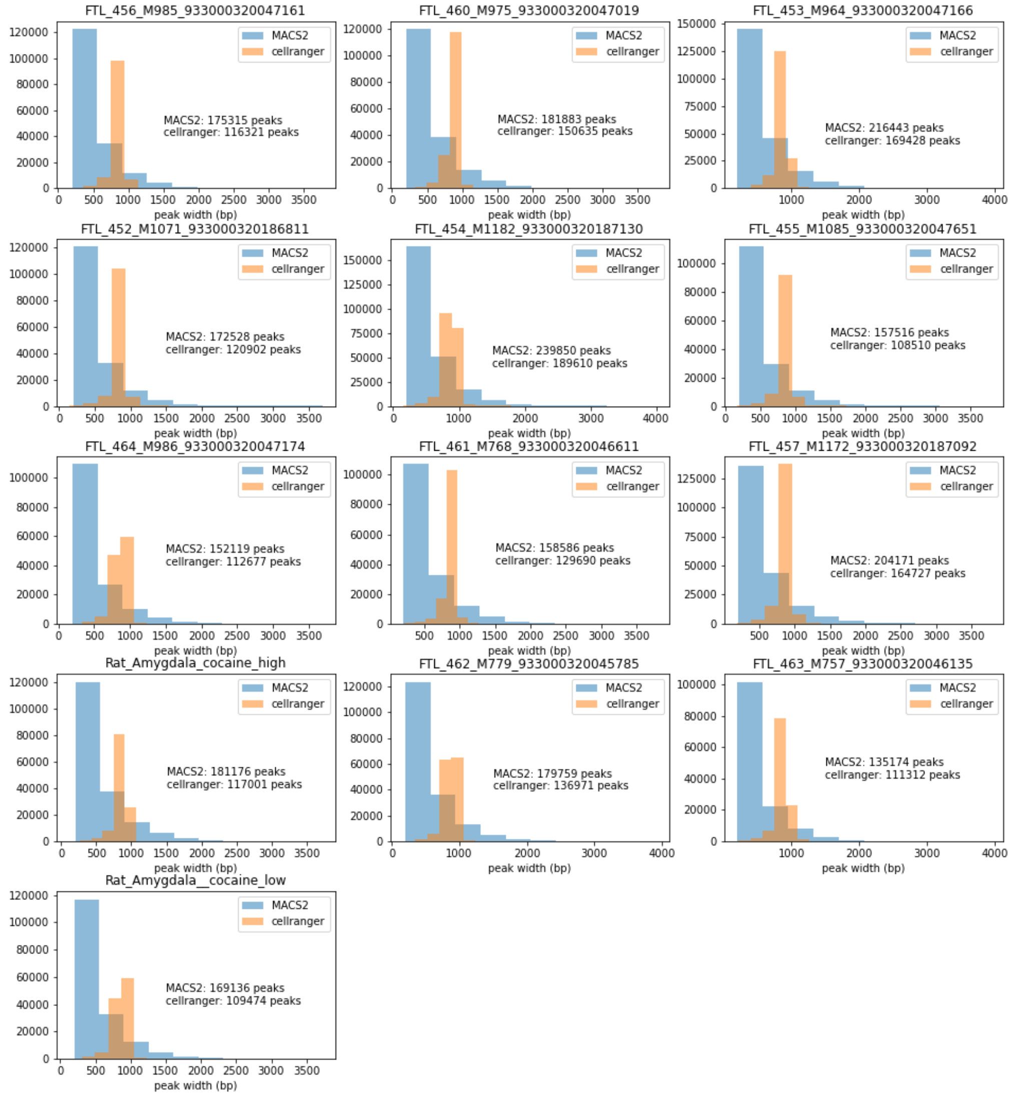


**Supplementary Figure 25.** Histograms showing distribution of peak sizes for peaks called by MACS2 (on the BAM files for the snATAC-seq data) versus CellRanger's internal peak calling algorithm. MACS2 calls smaller, more precise peaks.

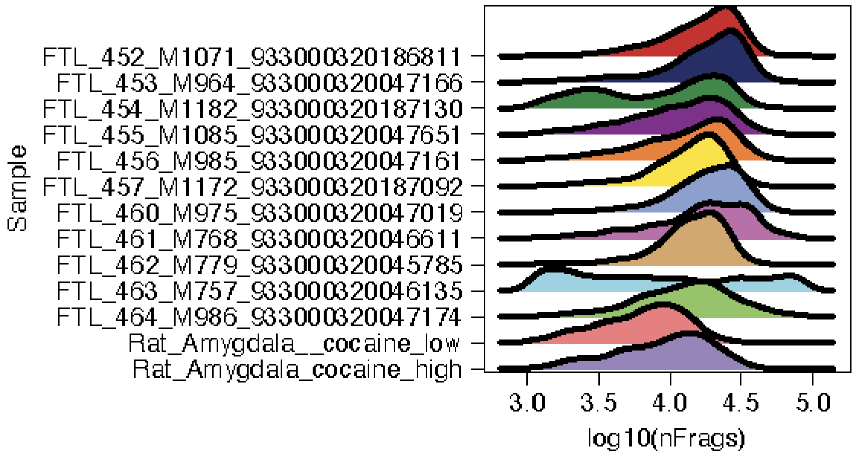


**Supplementary Figure 26.** Ridge plot quantifying the number of unique fragments (log10(nFrags)) per sample in the ATAC. Sample FTL_463_M757_933000320046135 was removed at this step and not included in any of our downstream snATAC-seq analyses due to its low number of fragments.

### Supplementary Tables

**Supplementary Table 1.** Number and percentage of nuclei per cell type for snRNA-seq and snATAC-seq.

| cluster | ncells.snRNA | percent.snRNA | ncells.snATAC | percent.snATAC |
|---|---|---|---|---|
| Astrocytes | 19651 | 12.05560634 | 7337 | 8.957173552 |
| Cck+/Vip+ | 3959 | 2.428789654 | 1496 | 1.82635023 |
| Chat+ | 1628 | 0.9987546241 | 1043 | 1.273317707 |
| Endothelial | 699 | 0.4288264633 | 949 | 1.158560406 |
| ExNeuron | 23943 | 14.68868671 | 20169 | 24.6227659 |
| InhNeuron | 52579 | 32.25646154 | 18208 | 22.22873327 |
| Microglia | 8834 | 5.419532156 | 4574 | 5.58404141 |
| Nos1+ | 4114 | 2.523879929 | 1232 | 1.50405313 |
| Oligodendrocytes | 29140 | 17.87697159 | 19391 | 23.67296611 |
| OPC | 9780 | 5.999889573 | 4031 | 4.921134876 |
| Pvalb+ | 423 | 0.2595044263 | 213 | 0.2600351597 |
| Reln+ | 1008 | 0.6183935265 | 426 | 0.5200703194 |
| Sst+ | 7245 | 4.444703472 | 2843 | 3.470797929 |

**Supplementary Table 2.** Results of Chi-squared test with Yates' continuity correction for enrichment of significant DEGs (FDR<10%) that also have eQTLs in the rat brain in each cell type.

| statistic | p.value | parameter | celltype | q.value |
|---|---|---|---|---|
| 32.00156145 | 1.54E-08 | 1 | Astrocytes | 9.24E-08 |
| 5.620791222 | 0.017748634 | 1 | Cck+-Vip+ | 0.050195845 |
| 263.0825833 | 3.65E-59 | 1 | ExNeuron | 3.29E-58 |
| 115.3577062 | 6.57E-27 | 1 | InhNeuron | 5.26E-26 |
| 8.589987545 | 0.003380163 | 1 | Microglia | 0.013520653 |
| 0.293805179 | 0.587792337 | 1 | Nos1+ | 0.587792337 |
| 30.98112175 | 2.61E-08 | 1 | Oligodendrocytes | 1.30E-07 |
| 5.724274495 | 0.016731948 | 1 | OPC | 0.050195845 |
| 43.28438553 | 4.73E-11 | 1 | Sst+ | 3.31E-10 |

**Supplementary Table 3.** Spearman correlations between difference in mean predicted expression and observed avg_logFC of expression between high vs. low AI rats for subsets of genes passing each Pearson $r^2$ cutoffs for gene expression prediction models.

| celltype | r2_cutoff | spearman_rho | pvalue | ci_low | ci_high | n_genes |
|---|---|---|---|---|---|---|
| Astrocytes | 0 | 0.018701122 | 0.283414263 | -0.015471691 | 0.052830298 | 3292 |
| Astrocytes | 0.025 | 0.018691166 | 0.357453182 | -0.021121508 | 0.058444654 | 2426 |
| Astrocytes | 0.05 | 0.02746218 | 0.250335653 | -0.019368071 | 0.074172197 | 1754 |
| Astrocytes | 0.075 | 0.036876648 | 0.170493112 | -0.015866717 | 0.08941536 | 1383 |
| Astrocytes | 0.1 | 0.039827961 | 0.180354767 | -0.018455348 | 0.097841507 | 1133 |
| ExNeuron | 0 | 0.025345141 | 0.145978786 | -0.00882543 | 0.059456588 | 3292 |
| ExNeuron | 0.025 | 0.034691131 | 0.087575896 | -0.00511289 | 0.074385397 | 2426 |
| ExNeuron | 0.05 | 0.042715472 | 0.073695635 | -0.004098078 | 0.089342205 | 1754 |
| ExNeuron | 0.075 | 0.048777171 | 0.069770271 | -0.003945496 | 0.101229416 | 1383 |
| ExNeuron | 0.1 | 0.049994674 | 0.092565284 | -0.008269905 | 0.107920939 | 1133 |
| InhNeuron | 0 | 0.033607128 | 0.053848494 | -0.000556439 | 0.067692338 | 3292 |
| InhNeuron | 0.025 | 0.046521106 | 0.021938511 | 0.006736615 | 0.086158555 | 2426 |
| InhNeuron | 0.05 | 0.064898807 | 0.006549113 | 0.018148579 | 0.111365876 | 1754 |
| InhNeuron | 0.075 | 0.057359357 | 0.03292901 | 0.004660927 | 0.109740074 | 1383 |
| InhNeuron | 0.1 | 0.064577247 | 0.0297394 | 0.00636067 | 0.122357559 | 1133 |
| Microglia | 0 | 0.035639761 | 0.040880673 | 0.001478633 | 0.069717804 | 3292 |
| Microglia | 0.025 | 0.060322746 | 0.002955403 | 0.020575177 | 0.099879931 | 2426 |
| Microglia | 0.05 | 0.062580891 | 0.008750828 | 0.015821879 | 0.109066777 | 1754 |
| Microglia | 0.075 | 0.061998527 | 0.021123002 | 0.009316451 | 0.114337383 | 1383 |
| Microglia | 0.1 | 0.05067189 | 0.08822582 | -0.007591011 | 0.108591918 | 1133 |
| OPC | 0 | 0.015433019 | 0.376048454 | -0.01873979 | 0.049569812 | 3292 |
| OPC | 0.025 | 0.01388068 | 0.494375796 | -0.025930607 | 0.053648007 | 2426 |
| OPC | 0.05 | 0.012821107 | 0.59154584 | -0.034004655 | 0.059590703 | 1754 |
| OPC | 0.075 | 0.012455566 | 0.64350377 | -0.040283405 | 0.065125329 | 1383 |
| OPC | 0.1 | -0.002824071 | 0.924351618 | -0.061054529 | 0.055425544 | 1133 |
| Oligodendrocytes | 0 | 0.062043491 | 0.000368203 | 0.027939783 | 0.096002933 | 3292 |
| Oligodendrocytes | 0.025 | 0.074118064 | 0.000258399 | 0.034422637 | 0.113580003 | 2426 |
| Oligodendrocytes | 0.05 | 0.084101822 | 0.000421907 | 0.037443861 | 0.130393894 | 1754 |
| Oligodendrocytes | 0.075 | 0.096983393 | 0.000303911 | 0.044498356 | 0.148934526 | 1383 |
| Oligodendrocytes | 0.1 | 0.102632954 | 0.000540258 | 0.044659402 | 0.159917563 | 1133 |

**Supplementary Table 4.** Results of two-sided Fisher's exact test measuring enrichment of DEGs with differentially accessible promoters.

| estimate | p.value | conf.low | conf.high | celltype | q.value |
|---|---|---|---|---|---|
| 18.81516404 | 0 | 17.77574265 | 19.9175995 | Astrocytes | 0 |
| 30.37291625 | 0.005840132 | 2.201548611 | 415.3908706 | Endothelial | 0.005840132 |
| 23.96758633 | 0 | 22.74043054 | 25.23646105 | ExNeuron | 0 |
| 17.57693406 | 0 | 16.48571015 | 18.74700536 | InhNeuron | 0 |
| 21.28725044 | 0 | 19.94126124 | 22.71783229 | Microglia | 0 |
| 27.72361083 | 0 | 26.31595904 | 29.21625885 | Oligodendrocytes | 0 |
| 10.28809974 | 7.95E-48 | 7.889775684 | 13.29876263 | OPC | 1.06E-47 |
| 30.37610297 | 0.000602346 | 4.068618798 | 226.6446137 | Pvalb+ | 0.000688396 |

**Supplementary Table 5.** Results of two-sided Fisher's exact test measuring enrichment of differentially accessible promoter regions (FDR<10%) in genes belonging to the oxidative phosphorylation pathway.

| estimate | p.value | conf.low | conf.high | celltype |
|---|---|---|---|---|
| 1.590213356 | 0.027413877 | 1.036306859 | 2.4171411 | Astrocytes |
| 0 | 1 | 0 | Inf | Cck+-Vip+ |
| 0 | 1 | 0 | Inf | Chat+ |
| 0 | 1 | 0 | 542.2633381 | Endothelial |
| 2.603860013 | 2.75E-06 | 1.705313547 | 4.02696631 | ExNeuron |
| 2.077830782 | 0.00082396 | 1.33980214 | 3.17681751 | InhNeuron |
| 1.50676579 | 0.06905433 | 0.945014063 | 2.350921065 | Microglia |
| 0 | 1 | 0 | 181.0640654 | Nos1+ |
| 3.044756686 | 6.43E-08 | 1.985216478 | 4.741095202 | Oligodendrocytes |
| 1.879741784 | 0.294615891 | 0.222019752 | 7.117370798 | OPC |
| 0 | 1 | 0 | 106.9419357 | Pvalb+ |
| 0 | 1 | 0 | Inf | Sst+ |

**Supplementary Table 6.** Results of two-sided Fisher's exact test measuring enrichment of differential peaks with TSS/promoter annotations.

| estimate | p.value | conf.low | conf.high | celltype | q.value |
|---|---|---|---|---|---|
| 18.81516404 | 0 | 17.77574265 | 19.9175995 | Astrocytes | 0 |
| 30.37291625 | 0.005840132 | 2.201548611 | 415.3908706 | Endothelial | 0.005840132 |
| 23.96758633 | 0 | 22.74043054 | 25.23646105 | ExNeuron | 0 |
| 17.57693406 | 0 | 16.48571015 | 18.74700536 | InhNeuron | 0 |
| 21.28725044 | 0 | 19.94126124 | 22.71783229 | Microglia | 0 |
| 27.72361083 | 0 | 26.31595904 | 29.21625885 | Oligodendrocytes | 0 |
| 10.28809974 | 7.95E-48 | 7.889775684 | 13.29876263 | OPC | 1.06E-47 |
| 30.37610297 | 0.000602346 | 4.068618798 | 226.6446137 | Pvalb+ | 0.000688396 |
