## Supplementary Data Guide for "Cocaine addiction-like behaviors are associated with long-term changes in gene regulation, energy metabolism, and GABAergic inhibition within the amygdala"

File name: Supplementary Data 1
Description: List of snRNA-seq rat samples included in analysis, their addiction indexes, batch information, and Cell Ranger summary metrics.

File name: Supplementary Data 2
Description: List of snATAC-seq rat samples included in analysis, their addiction indexes, batch information, and Cell Ranger summary metrics.

File name: Supplementary Data3
Description: Per-nucleus metrics for all nuclei in snRNA-seq dataset after filtering. This table contains selected columns from the metadata table for the Seurat object containing the integrated snRNA-seq data.

File name: Supplementary Data 4
Description: Per-nucleus metrics for all nuclei in snATAC-seq dataset after filtering. This table contains selected columns from the metadata table for the Signac object containing the integrated snATAC-seq data.

File name: Supplementary Data 5
Description: All cell type-specific differential gene expression analysis results, obtained using the negative binomial test.

File name: Supplementary Data 6
Description: Results of permutation test for differential gene expression analysis using negative binomial test.

File name: Supplementary Data 7
Description: All DEGs (FDR<10%) that also have eQTLs in the rat brain, with a list of variant IDs for corresponding eQTLs.

File name: Supplementary Data 8
Description: KEGG GSEA results.

File name: Supplementary Data 9
Description: All cell type-specific differential peak accessibility analysis results, obtained using the negative binomial test.

File name: Supplementary Data 10
Description: Permutation test for differential peak accessibility analysis results using negative binomial test.

File name: Supplementary Data 11
Description: ChromVar analysis results.
